# Chemically-induced targeted protein degradation in mycobacteria uncovers antibacterial effects and potentiates antibiotic efficacy

**DOI:** 10.1101/2023.02.14.528552

**Authors:** Harim I. Won, Junhao Zhu, Olga Kandror, Tatos Akopian, Ian D. Wolf, Michael C. Chao, Maya Waldor, Eric J. Rubin

## Abstract

Proteolysis-targeting chimeras (PROTACs) represent a new therapeutic modality involving selectively directing disease-causing proteins for degradation through proteolytic systems. Our ability to exploit this targeted protein degradation (TPD) approach for antibiotic development remains nascent due to our limited understanding of which bacterial proteins will be labile TPD targets. Here, we use a genetic system to model chemically-induced proximity and degradation to screen essential proteins in *Mycobacterium smegmatis* (*Msm*), a model for the major human pathogen *M. tuberculosis* (*Mtb*). We find that drug-induced proximity to the bacterial ClpC1P1P2 proteolytic complex is sufficient to degrade many, but not all, endogenous *Msm* proteins, profoundly inhibiting bacterial growth for some targets. We also show that TPD can potentiate the effects of existing antibiotics targeting the same pathways and complexes. Together, our results identify specific endogenous mycobacterial proteins as attractive targets for future *Mtb* PROTAC development, as both standalone antibiotics and potentiators of existing antibiotic efficacy.

## Introduction

Antibiotics, like many small molecule therapeutics, traditionally exert their mechanism of action through modulating a specific molecular target, whether by direct binding to an enzymatic active site or allosteric conformational changes. Efforts to develop new antibiotics largely apply this same paradigm, but development remains challenging due to the extremely high failure rate^1^ and high cost^2^. As a result, the discovery of new antibiotics has stalled dramatically in recent decades, including of those to treat bacterial infections like tuberculosis (TB), which remains one of the world’s leading infectious killers. The continued use of old antibiotics has compounded this problem, by giving rise to multi-drug resistant TB strains which are incredibly difficult to treat^3^. Because the traditional approach of single target modulation has slowed in its ability to discovery powerful new anti-tubercular agents, we sought to apply a novel modality to TB antibiotic development.

Targeted protein degradation (TPD) is an emerging therapeutic modality that has progressed from concept^4^ to Phase 2 clinical trials^5^ within the past two decades. The major class of molecules that enable protein level regulation through TPD are the proteolysis-targeting chimeras (PROTACs). These molecules are heterobifunctional, comprising two small molecule ligands joined by a chemical linker. One of the ligands binds an E3 ubiquitin ligase while the other binds a given target protein. The PROTAC induces the proximity of both the target protein and the E3 ligase, resulting in polyubiquitination of the target and its subsequent degradation by the eukaryotic proteasome. Other TPD approaches employ the lysosomal^6^ and autophagic^7,8^ degradation machinery.

Despite its great potential, the development of TPD-based antimicrobials remains largely conceptual. Rather than using host degradation machinery in a TPD strategy for bacteria, induced autoproteolysis would instead deliver proteins essential for bacterial viability to bacterial degradation machinery, resulting in their degradation and subsequent cell death or growth inhibition. In mycobacteria, options for degradation machinery include the mycobacterial proteasome, which uses the PafA ligase (a system analogous to the ubiquitination system employed by PROTACs^9^), or the Clp proteolytic complexes (i.e., ClpC1P1P2 and ClpXP1P2, which are essential in mycobacteria).

Recently, the first TPD proof-of-concept in bacteria demonstrated that heterobifunctional bacterial PROTACs (BacPROTACs) could mediate the inducible degradation of target proteins in *Msm* cells^10^. This approach relied on direct delivery of target proteins to the multicomponent, ClpC1P1P2 proteolytic complex (Fig. 1a), resulting in their degradation. In this work, BacPROTAC-mediated degradation of different targets was shown to induce sensitivity to D-cycloserine or auxotrophy for L-threonine, highlighting the potential of TPD as the foundation of a new class of antibiotic degraders. However, it remains unclear whether all proteins are equally good targets for this approach, and whether some are more attractive targets for future drug development. To determine this, we developed a rational platform for TPD target selection in mycobacteria, where we surveyed a set of diverse, essential mycobacterial proteins to examine their suitability for a TPD approach. Our genetic platform enables the rapid assessment of different target proteins for TPD, identifying suitable substrates for degradation. We also find that targeted degradation of particular substrates is sufficient to elicit cellular growth inhibition and potentiation of killing by clinical TB antibiotics even when targeted degradation alone functions at sub-inhibitory levels.

**Fig. 1.**
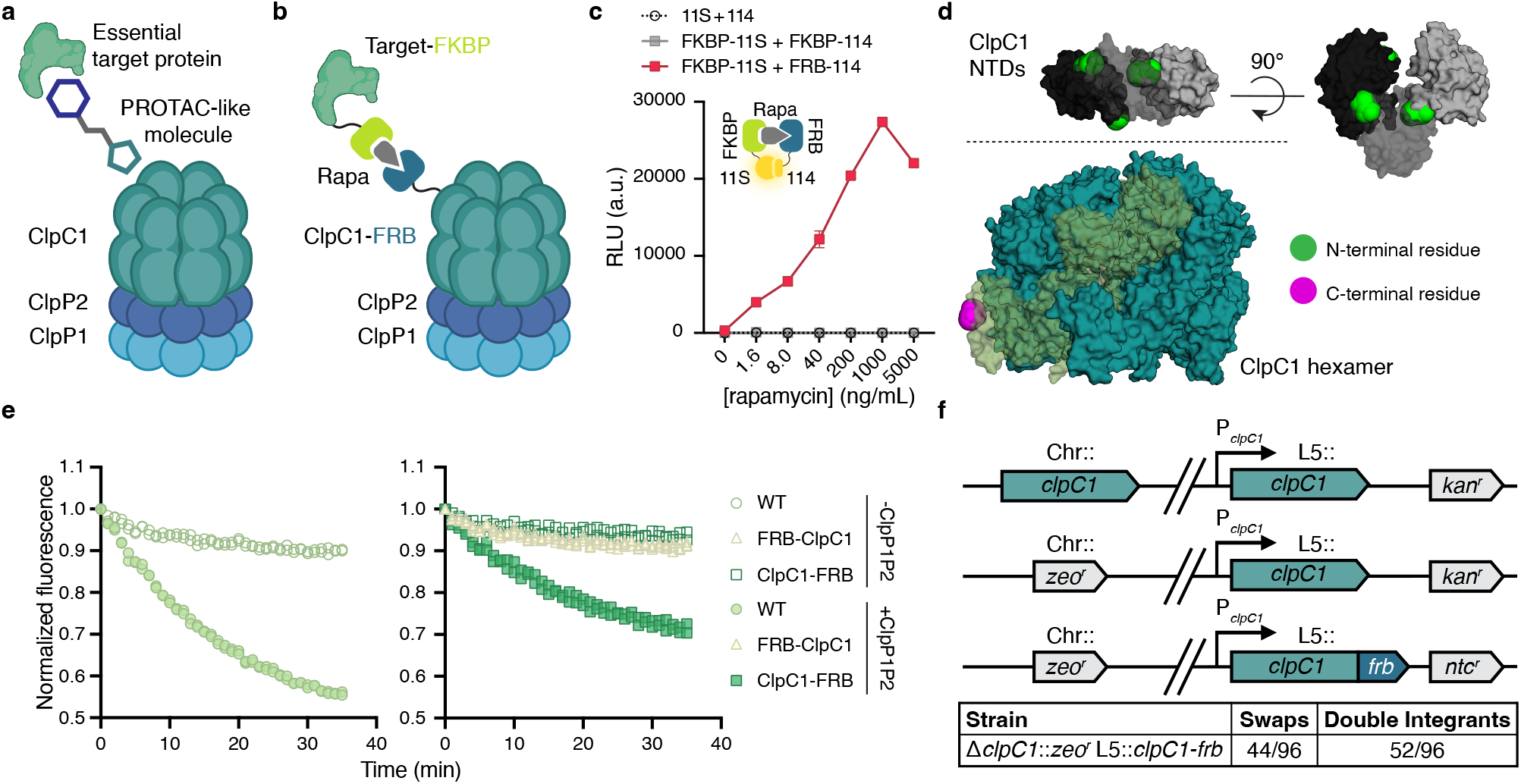
The FRB-FKBP dimerizable domains can be used to induce proximity in mycobacteria. **a** The concept of bacterial targeted protein degradation (TPD), in which an essential target protein is delivered to the ClpC1P1P2 proteolytic complex by a heterobifunctional molecule. **b** Schematic of our chemical genetic approach to induced proximity, in which fusions to the FRB and FKBP domains are dimerized with the addition of rapamycin. **c** Luminescence in live cells expressing split NanoLuc (Nluc) fragments with and without fusion to FRB and FKBP. Density matched log phase cells incubated with the luciferase substrate furimazine and a range of rapamycin concentrations at 37°C for 10 min. **d** Crystal structure of the stabilized mutant *Mtb*ClpC1 hexamer (teal, PDB: 8A8U) with highlighted monomer (translucent green) and three visible N-terminal domains (NTDs) which cannot be assigned to specific protomers due to invisibility of the linker region (grayscale, PDB: 6PBS). ClpC1 N- and C-terminal residues highlighted in green and magenta, respectively. **e** Fluorescence of eGFP-ssrA measuring *in vitro* protease activity of ClpC1 P1P2 complex with WT *Mtb*ClpC1 (left) or FRB-ClpC1 and ClpC1-FRB (right). Purified *Mtb*Clp proteins incubated with eGFP-ssrA substrate at 37°C. **f** (Top) Schematic of the *clpC1* L5 allele swap, in which a second copy of *clpC1* is integrated at the L5 phage *attB* site. This enables recombineering-mediated knockout of chromosomal *clpC1*, followed by an L5 integrase-mediated swap for the *clpC1-frb* allele and an alternative resistance marker. (Bottom) Quantification of the *clpC1-frb* swap; true swaps carry only the second resistance marker, whereas double integrants carry both (and both *clpC1* alleles). For **c**, data are mean ± s.d. of three technical replicates and are representative of two independent experiments; **e**, data are individually plotted technical replicate measurements, normalized to time = 0 h, and are representative of two independent experiments. **a-b** Created with BioRender.com

## Results

### A chemical genetic platform for evaluating targeted protein degradation in mycobacteria

The development of a targeted protein degradation (TPD) strategy for *Mtb* infection necessitates the identification of protein targets that are required for bacterial growth and are amenable to proximity mediated degradation by bacterial proteolytic systems. However, the systematic evaluation of vulnerable protein targets for TPD drug development is infeasible using a chemical-forward approach (i.e., identifying chemical ligands to all 461 genes identified as essential for *Mtb* growth^11^). Therefore, we sought to develop a platform for rational target selection, which would enable the prioritization of targets before chemical screening for binding ligands. To accomplish this, we needed a system that would enable regulated, inducible proximity of native proteins and a mycobacterial proteolytic complex like ClpC1P1P2, which has recently been used for TPD in bacteria^10^. For this purpose, we chose to create genetic fusions with FRB, the rapamycin-binding domain of mTOR, and FKBP12, which binds both rapamycin and FK506^12^; FRB and FKBP are small (~12 kDa) proteins which dimerize with nanomolar affinity only in the presence of rapamycin^13,14^. We fused ClpC1 with the FRB domain and candidate proteins with FKBP, creating strains where the presence of rapamycin induces the proximity of target proteins to ClpC1 (Fig. 1b). To enable rapid screening of targets which were most effectively degraded, we built this system in the non-pathogenic model of *Mtb, Mycobacterium smegmatis* (*Msm*).

Though FRB and FKBP have been widely used to conditionally dimerize proteins in mammalian cells^14^ and, more recently, in *Escherichia coli*^15^, it had not yet been evaluated in live mycobacterial cells. Therefore, we first assessed whether rapamycin-dependent dimerization occurred in *Msm* cells by fusing each domain to the NanoLuc (Nluc) 11S fragment and 114 peptide split reporter system^16^. Compared to other split reporter systems (e.g., fluorescent proteins, β-galactosidase), the split Nluc (NanoBiT) system features the rapid signal detection, sensitivity, and dynamic range of luciferase assays while also being optimized to have minimal affinity between the split components. We expressed the FRB and FKBP Nluc fusion proteins in *Msm* and observed rapamycin-dependent luciferase activity only when both rapamycin-binding domains were fused to the split Nluc fragments (i.e., FRB-11s and FKBP-114), and this activity appeared to saturate at a rapamycin concentration of 1 μg ml^-1^ (Fig. 1c). We also observed the characteristic “hook effect” resulting from the formation of unproductive binary complexes at high concentrations rather than necessary ternary complexes required for induced proximity^17,18^. Importantly, all concentrations of rapamycin we used to induce proximity in this work were inert with respect to bacterial growth and were orders of magnitude below the highest tested concentration which itself also did not inhibit bacterial growth in wild-type mc^2^155 strain *Msm* cells (Supplementary Figure 1).

To generate the ClpC1 and FRB fusion protein, the FRB domain could be fused to either the N-terminal or C-terminal end. Based on the crystal structures of the *Staphylococcus aureus* ClpC and *Mtb* ClpC1 hexamer^19,20^, we reasoned while N-terminal fusion might facilitate the direct delivery of target proteins to the ClpC1 hexamer pore, the genetic engineering of the ClpC1 C-terminus would be better tolerated by cells given the degree to which the N-terminal ends were nestled among the ClpC1 N-terminal domains (NTDs) compared to the relative accessibility of the C-terminal end (Fig. 1d). To compare these two scenarios, we purified both the N-terminal and C-terminal *Mtb*ClpC1 fusions (FRB-ClpC1 and ClpC1-FRB, respectively; Supplementary Figure 2a) to measure each fusion protein’s ability to degrade the eGFP-ssrA target protein. This target was selected as WT *Mtb*ClpC1 in association with ClpP1P2 had been previously shown to be able to degrade eGFP-ssrA *in vitro^21,22^*. Indeed, we found FRB-ClpC1 had no measurable protease activity, while ClpC1-FRB was able to degrade eGFP-ssrA, albeit to a lesser degree than WT ClpC1 (Fig. 1e). We also observed this trend among the different ClpC1 fusion variants without ClpP1P2 when measuring ATPase activity, though here, FRB-ClpC1 showed some minimal activity (Supplementary Figures 2b, c).

To generate a strain carrying a single copy of *clpC1-frb* under the native *clpC1* promoter, we replaced wild-type *clpC1* with a *clpC1-frb* allele via an L5 integrase allele swap^23^ (Fig. 1f, top). We generated viable *Msm* strains with clean swaps for *clpC1-frb* alleles at a rate of 45.8%, successfully verifying that ClpC1-FRB remains functional *in vivo* and can complement wild-type ClpC1’s essential role in normal mycobacterial growth (Fig. 1f, bottom). We further confirmed that the addition of the FRB domain did not disrupt cellular physiology by measuring growth and found that the strain expressing ClpC1-FRB shared growth characteristics with parental *Msm* (Supplementary Figure 3). This strain served as the basis for our platform to assess TPD in mycobacteria and examine whether we could mediate the degradation of specific target proteins by induced proximity to ClpC1.

### Rapamycin-induced proximity to ClpC1 is sufficient to degrade a native mycobacterial protein

To test whether rapamycin-induced proximity to ClpC1 was sufficient for mycobacterial protein degradation, we constructed target protein fusions with FKBP and enhanced green fluorescent protein (eGFP) and introduced them as merodiploid copies into the Tweety phage *attB* site^24^ in both a WT *clpC1* or *ΔclpC1 L5::clpC1-frb* background (Figs. 2a, b) and measured the effect of rapamycin on fluorescence in *Msm*. Before testing different native targets, we assessed whether the minimal backbone, FKBP-eGFP alone, could be directed for degradation by ClpC1 or if degradation was more substrate selective. We found that heterologously expressed FKBP-eGFP was not degraded regardless of *clpC1* background (Fig. 2c), suggesting either that it is a poor ClpC1 substrate (perhaps due to eGFP’s characteristic, protease-resistant tight folding^25^) or that the geometry of delivery we achieved is incompatible with efficient degradation. To test whether the absence of rapamycin-induced degradation was due to possible cryptic obstruction of the rapamycin-mediated protein-protein interaction, we examined the subcellular distribution of the FKBP-eGFP signal with or without rapamycin treatment. Previous work indicates that full-length ClpC1-eGFP locates near the plasma membrane and enriches at the subpolar region in a non-homogeneous manner^26^ which we also observed (Supplementary Figures 4a, b). While FKBP-eGFP in the *clpC1-frb* background diffuses in the cytosol, the addition of rapamycin results in re-localization of FKBP-eGFP to the subpolar membrane compartment (Supplementary Figure 4c), indicating that rapamycin successfully mediates the physical interaction between FKBP-eGFP and ClpC1-FRB and further suggesting that FKBP-GFP alone is a suboptimal substrate for ClpC1.

**Fig. 2.**
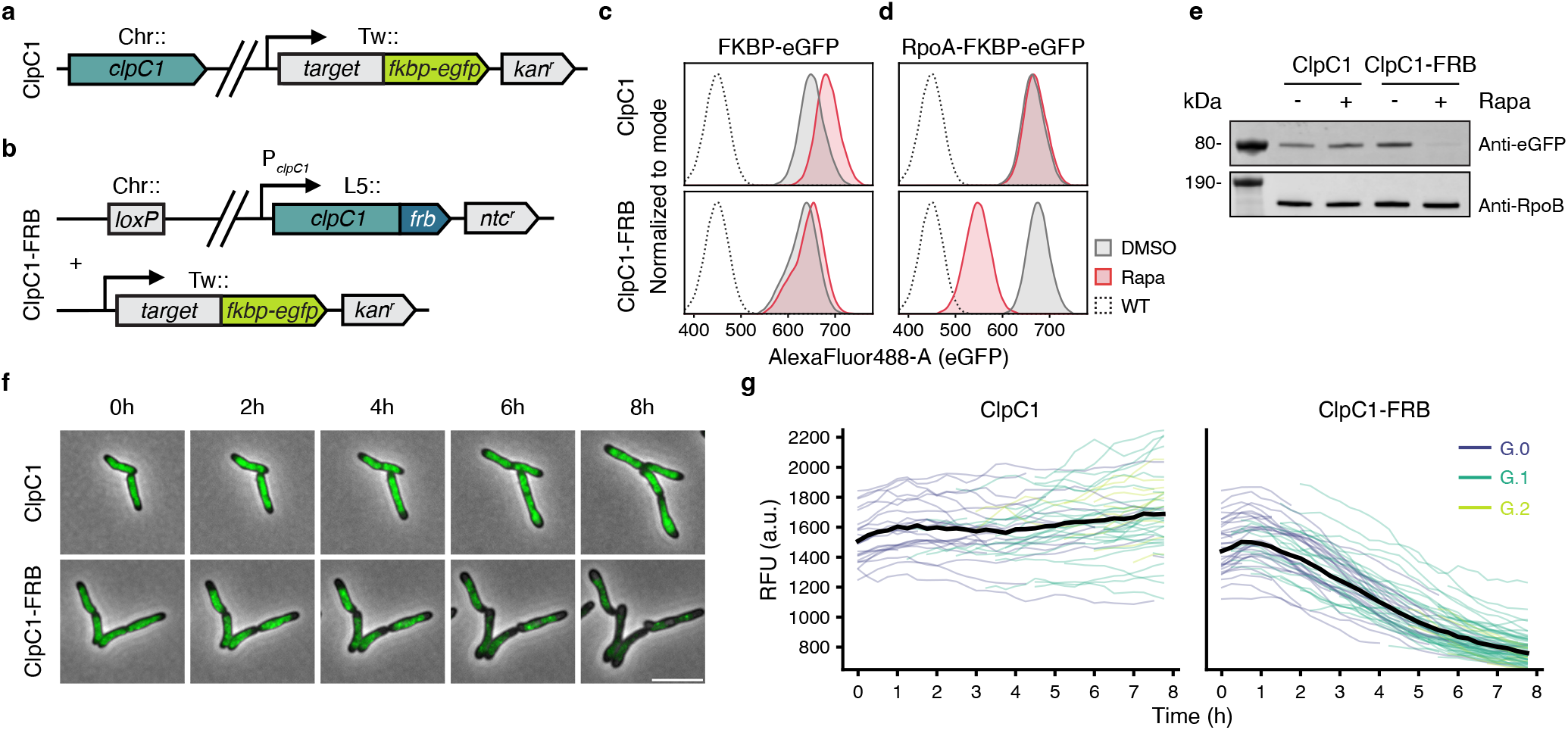
Rapamycin redirects a native mycobacterial protein for degradation by ClpC1-FRB. **a-b** Schematics of reporter strains generated to assess target degradation, with *substrate-fkbp-egfp* genes delivered to the Tweety phage *attB* site in a WT *clpC1* (**a**) or *ΔclpC1 L5::clpC1-frb* (**b**) background. **c-d** Fluorescence of live cells as a proxy for protein levels of FKBP-eGFP (**c**) or RpoA-FKBP-eGFP (**d**) in the WT *clpC1* (top) or *clpC1-frb* (bottom) background. Density matched log phase cells incubated with DMSO or 0.1 μg ml^-1^ rapamycin with shaking at 37°C for 24 h. **e** Western blot analysis of RpoA-FKBP-eGFP with DMSO or 0.1 μg ml^-1^ rapamycin addition in both strain backgrounds. **f** Live cell, wide-field fluorescence microscopy time-lapse images of cells expressing RpoA-FKBP-eGFP with 0.1 μg ml^-1^ rapamycin in both strain backgrounds. Scale bar, 5 μm. **g** Quantitation of all captured fields during time-lapse in (**f**), measuring median fluorescent signal in the FITC channel across cells over time. Colored lines represent individual cells (ClpC1, n=77; ClpC1-FRB, n=105) that were initially plated (G.0, purple), first-generation (G.1, teal), or second-generation (G.2, lime green). For **c-d**, data are representative of three independent experiments; **e**, data represent one independent experiment; **f-g**, data are representative images selected from among 4 fields for each and are representative of two independent experiments.

We next selected RpoA, a small (38 kDa) and essential component of the RNA polymerase (RNAP) holoenzyme, to test whether the addition of an endogenous protein could potentiate proximity-induced degradation by ClpC1. Indeed, we found that rapamycin is sufficient to direct RpoA-FKBP-eGFP for degradation, with substantial loss of eGFP signal only in the *clpC1-frb* background (Fig. 2d). To validate that the observed loss of eGFP signal was the result of *bona fide* protein degradation, we showed a rapamycin-dependent reduction in protein levels of the full length RpoA-FKBP-eGFP fusion protein only in the *clpC1-frb* background (Fig. 2e). Furthermore, to assess the dynamics of this targeted protein degradation of RpoA, we examined individual cells by time-lapse fluorescence microscopy and observed a rapamycin-dependent reduction of eGFP signal only in the *clpC1-frb* background within 8 hours of induction (Figs. 2f, g). Our findings demonstrate that a genetically encoded system for chemically-induced proximity to ClpC1 which is orthogonal to the chemical approach developed by Morreale *et al.^10^*, is sufficient to degrade a native mycobacterial protein.

### Mycobacterial proteins are differentially susceptible to ClpC1-mediated TPD

To identify suitable endogenous targets for TPD more systematically, we fused a set of native mycobacterial proteins to FKBP-eGFP and assessed their degradation using high-throughput flow cytometry (Fig. 3a and Supplementary Figure 6a).

**Fig. 3.**
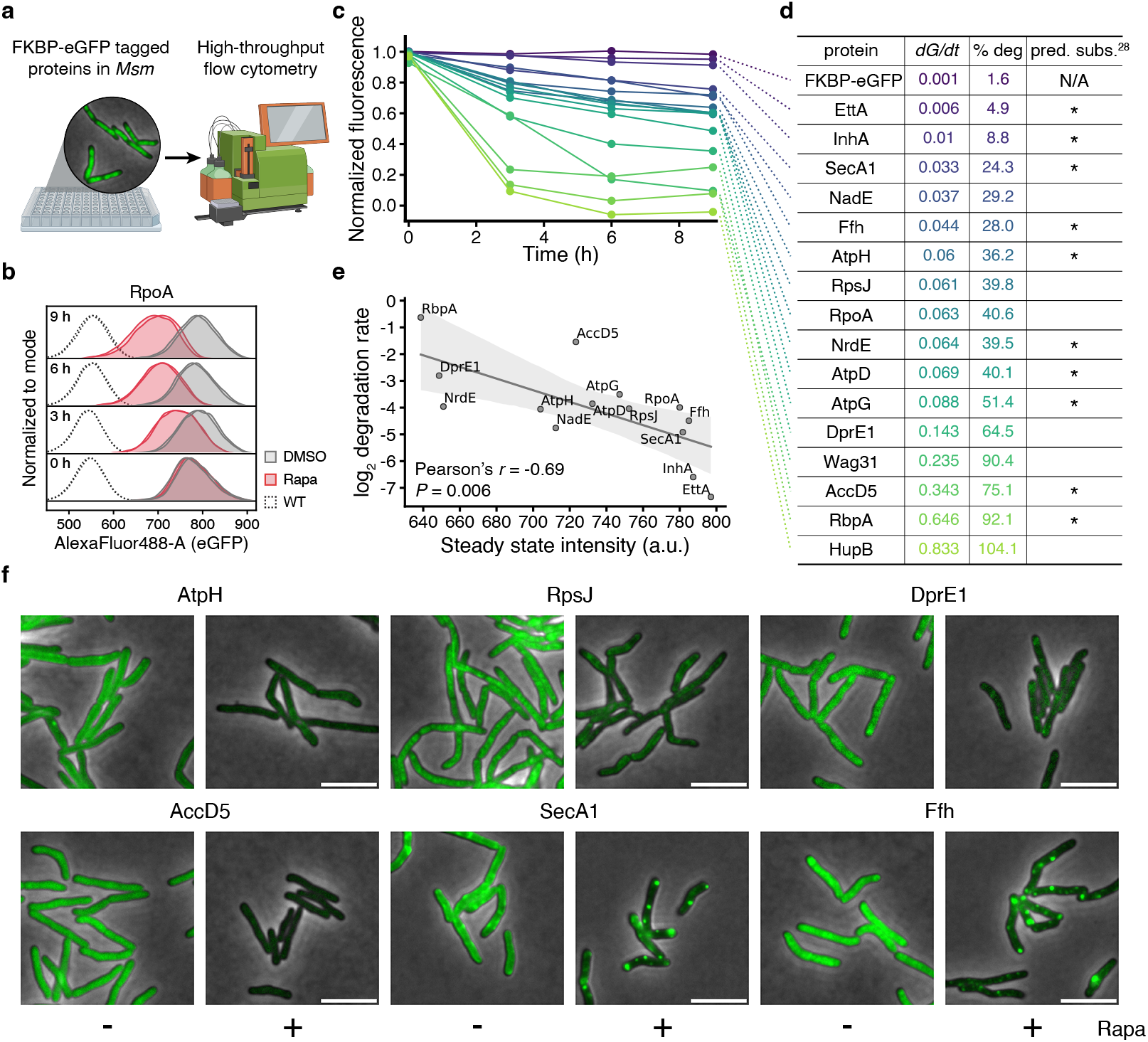
Mycobacterial proteins are differentially susceptible to degradation by ClpC1-FRB. **a** Schematic of high-throughput flow cytometry screen in *Msm* cells expressing targets tagged with FKBP-eGFP. **b** Fluorescence of live cells as a proxy for protein levels of RpoA-FKBP-eGFP in the *clpC1-frb* background over time. **c** Summary plot of all tested target proteins; fluorescence of live cells as a proxy for protein levels in the *clpC1-frb* background over time. Density matched log phase cells incubated with DMSO or 0.5 μg ml^-1^ rapamycin with shaking at 37°C for the indicated times. **d** Enumeration of all tested proteins and their respective rate constants (*dG/dt*), proportion of total degradation (% deg), and whether they are predicted to be native ClpC1 substrates^28^. **e** Correlation plot relating the log2-transformed degradation rate and the steady state intensity of plotted target proteins. **f** Live cell, wide-field fluorescence microscopy images of cells expressing selected target proteins with DMSO or 0.1 μg ml^-1^ rapamycin. Scale bar, 5 μm. For **b**, data are two technical replicates, are representative of three independent experiments, and normalized to the mode; **c**, data are the mean of two technical replicates, are representative of two independent experiments, and normalized to fluorescent intensity of DMSO-treated cells at time = 0 h. Data in **e** are bounded by the 95% confidence interval. Data in **f** are representative images selected from among 4 fields for each and are representative of two independent experiments. **a** Created with BioRender.com

The fluorescent degradation kinetics of RpoA as measured by flow cytometry (Fig. 3b) correlated well with the signal observed by time-lapse microscopy (Supplementary Figure 6b), motivating our use of flow cytometry as the basis for our target screening assay. We selected additional essential targets which were components of central biological processes (e.g., replication, translation, ATP synthesis) and had distinct subcellular localizations^27^ (to assess potential challenges with re-localizing certain targets to ClpC1). We quantified the effect of rapamycin on the fluorescence of a total of 17 targets over time and found that most of the tested native proteins were able to be targeted for degradation by rapamycin (Fig. 3c and Supplementary Figure 7a) and that the observed fluorescent signal decay fit well to an exponential decay function (Supplementary Figure 7b), permitting us to calculate a rate constant for each target (Fig. 3d).

While some targets display slower degradation kinetics – such as RpoA, which did not reach maximal, steady state degradation until 9 hours (*dG/dt* = 0.063) – we found that others, such as the transcription factor RbpA, were rapidly degraded to a steady state baseline before 3 hours (*dG/dt* = 0.646; Fig. 3d and Supplementary Figure 7a). Intriguingly, efficient TPD by ClpC1 was not restricted to native ClpC1 substrates, as we compared our list of proteins to a proteomics dataset that predicted a set of native ClpC1 substrates^28^ and found no correlation between being a native substrate and degradation efficiency (Fig. 3d).

Interestingly, even though all targets were expressed from the same constitutive promoter (except Wag31 and HupB, due to toxicity), we observed variance in the steady state fluorescence in the different strains in the absence of rapamycin, indicating that there may be some basal post-translational regulation of some targets, possibly through proteolysis. We wondered whether these “active degraders” might be more susceptible to TPD, perhaps because these proteins had exposed degrons that made them more amenable to protease activity. Indeed, we found an inverse correlation with the log2 degradation rate of each target and its steady state fluorescence (Fig. 3e; Pearson’s *r* = −0.69, *P* = 0.006).

To assess the effect of rapamycin-induced proximity to ClpC1 on protein localization and protein levels, we imaged each strain using fluorescent, live-cell microscopy (Fig. 3f and Supplementary Figures 8a, b). We confirmed the gradient of degradability of the tested targets and again found high concordance between the flow cytometry and microscopy datasets (Supplementary Figure 8c). Noticeably, for SecA1 and Ffh, which are only moderately degraded by TPD, the addition of rapamycin resulted in profound re-localization of fluorescent signal (Fig. 3f). Though these observations could reflect biology, these punctate foci could alternatively form due to oligomerization of eGFP when local concentrations increase near ClpC1-FRB. Rapamycin does appear to be stable in this system as eGFP levels remain low for at least 48 h post-inoculation (Supplementary Figure 9).

### Targeted protein degradation of native mycobacterial proteins inhibits bacterial growth

In our previous experiments revealing that induced proximity drives the degradation of several mycobacterial proteins, we did not expect to see any impacts on growth because the strains we generated for these experiments were merodiploid for the protein being degraded (i.e., the genes were present in two copies). To examine the effect of targeted degradation of native mycobacterial proteins on bacterial growth and viability, we endogenously tagged the various targets at their chromosomal loci with *fkbp* by recombineering in both a WT *clpC1* or *ΔclpC1 L5::clpC1-frb* background (Figs. 4a, b) and measured the effect of rapamycin on bacterial growth. We observed that rapamycin-mediated degradation of RpoA and the ATP synthase component AtpA resulted in a mild growth delay in cells grown in liquid media only in the *clpC1-frb* background (Supplementary Figure 10). In contrast, we observed strong growth delays when cells targeting AtpA and Ffh for degradation were plated on media containing rapamycin (Fig. 4c) and further demonstrated that degradation of Ffh and AtpA resulted in the formation of ~24% fewer colonies (Fig. 4d; AtpA *P* = 0.0083, Ffh *P* = 0.0169). Quantitative plate image analysis also confirmed delayed colony outgrowth when degrading AtpA and Ffh, only in the *clpC1-frb* background (Fig. 4e and Supplementary Figure 11). It is worth noting that Ffh is a component of the bacterial version of the signal recognition particle (SRP) system, which could mean that effective depletion of this target might additionally disrupt the proper secretion of essential membrane proteins. Interestingly, targeted degradation of other components of the ATP synthase machinery (i.e., AtpH, AtpG, AtpD) did not manifest the outgrowth delay we observed with AtpA. Together, these results suggest that efficient degradation of a given target is necessary, but not sufficient for growth phenotypes *in vivo;* therefore, selection of a target must be validated empirically at the chromosomal locus even after effective degradation is verified.

**Fig. 4.**
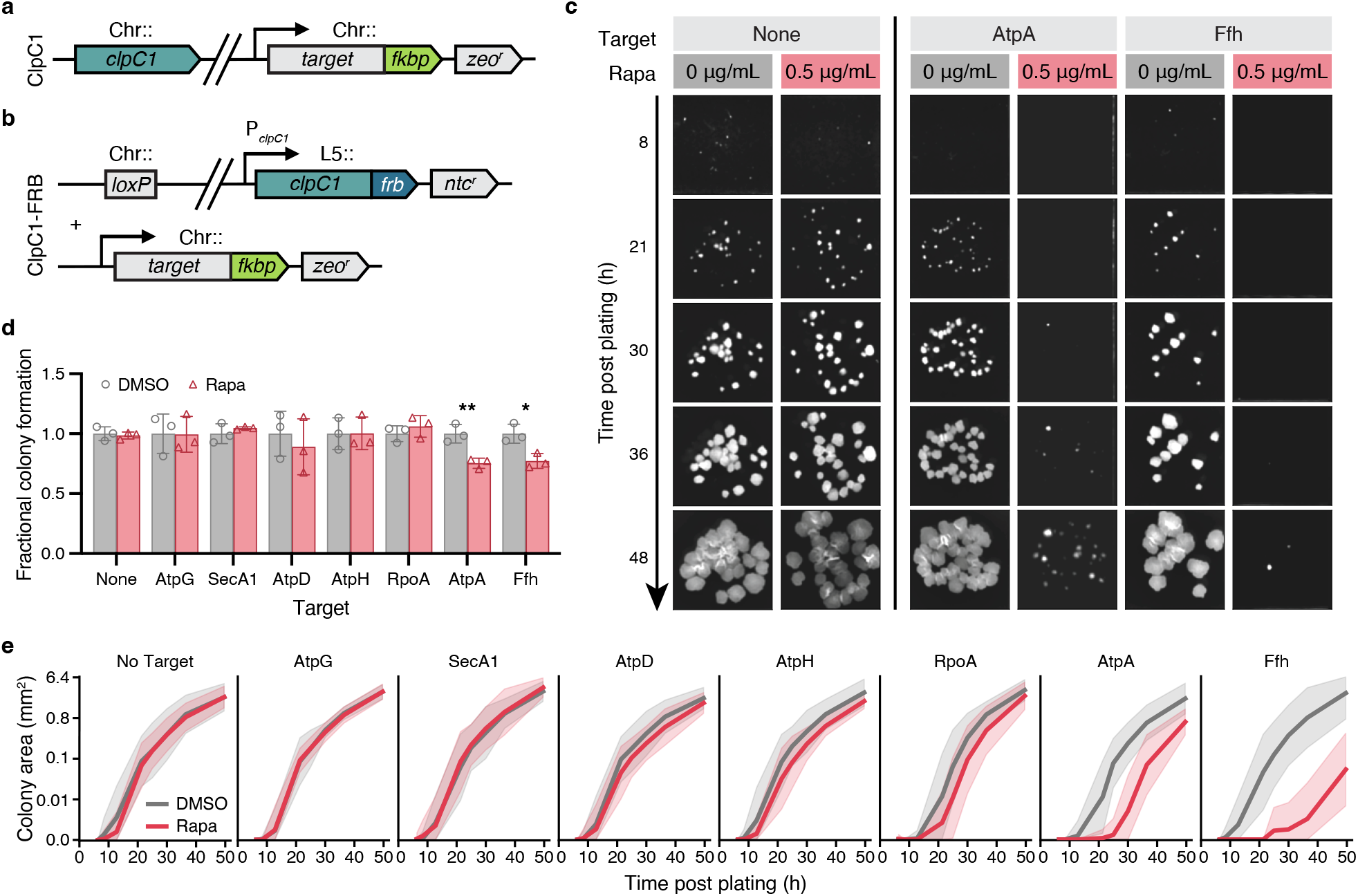
Targeted degradation of mycobacterial proteins inhibits bacterial growth. **a-b** Schematics of strains generated to assess phenotypes arising from targeted protein degradation, with target alleles tagged with *fkbp-flag (atpG, secA1, atpD, atpH, rpoA, atpA*) or *fkbp-egfp (ffh*) at their chromosomal loci in a WT *clpC1* (**a**) or *ΔclpC1 L5::clpC1-frb* (**b**) background. **c** Representative images illustrating colony outgrowth dynamics of the indicated targets in the *clpC1-frb* background over time. Density matched log phase cells serially diluted, plated on solid media containing DMSO or 0.5 μg ml^-1^ rapamycin, and incubated at 37°C for the indicated times. **d** Total colonies formed during outgrowth on solid media containing DMSO or 0.5 μg ml^-1^ rapamycin for the indicated targets in the *clpC1-frb* background. *P* values were determined by unpaired two-tailed *t*-tests and compared 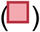 with 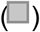. \**P* < 0.05, \*\**P* < 0.01. Exact *P*-values: AtpA, \*\**P* = 0.0083; Ffh, \**P* = 0.0169. **e** Quantitation of colony outgrowth dynamics as in (**c**) by colony size tracking by area (mm^2^) of individual colonies of the indicated targets in the *clpC1-frb* background over time. Cells plated as in **c-d**. For **c**, images are representative of three technical triplicates and two independent experiments; **d**, data are mean ± s.d. of three technical replicates and are representative of two independent experiments. In **e**, dark lines are the mean of three technical replicates, are bounded by the 95% confidence interval, and are representative of two independent experiments.

### Targeted degradation of druggable complexes sensitizes mycobacteria to corresponding antibiotics

In addition to the growth phenotypes we observed above, we examined whether targeted degradation of protein complex components might sensitize bacteria to clinically relevant antibiotics that target different members of those complexes. Specifically, we assessed whether targeted degradation of RpoA would synergize with rifampicin, which targets the RpoB subunit of the RNA polymerase holoenzyme; and if targeted degradation of AtpA would synergize with bedaquiline, which targets the AtpE subunit of ATP synthase (Fig. 5a). In both cases, with targeted degradation of RpoA and AtpA, cells were more sensitive to rifampicin and bedaquiline, respectively (Figs. 5b, c). These observed shifts represented a 4.13-fold shift for the MIC_50_ of rifampicin and a 3.67-fold shift for the MIC_50_ of bedaquiline. There was no sensitization to either antibiotic in cells where targeted degradation was not occurring (Supplementary Figures 12a, b). We also examined whether RpoA and AtpA degradation sensitized cells to the antibiotics streptomycin and linezolid which inhibit translation by targeting the 30S and 50S ribosomal subunits, respectively. There was no sensitization to streptomycin when degrading RpoA, but we did observe a 2.69-fold shift in the MIC_50_ of streptomycin when degrading AtpA (Supplementary Figure 12c). Degrading both RpoA and AtpA resulted in 1.48-fold and 2.17-fold shifts in the MIC_50_ of linezolid (Supplementary Figure 12d). It is worth noting that the effect of degrading RpoA and AtpA in the presence of antibiotics targeting different pathways were less pronounced compared to antibiotics targeting the same enzyme complexes. We hypothesize that the observed slight, non-specific sensitization may be the result of cells being in a compromised state due to degradation of transcription or ATP synthesis machinery; while we expect these cells to be more susceptible to antibiotics targeting the same pathway that is being degraded, it is also possible that their compromised state also renders them weaker against antibiotic stress more generally.

**Fig. 5.**
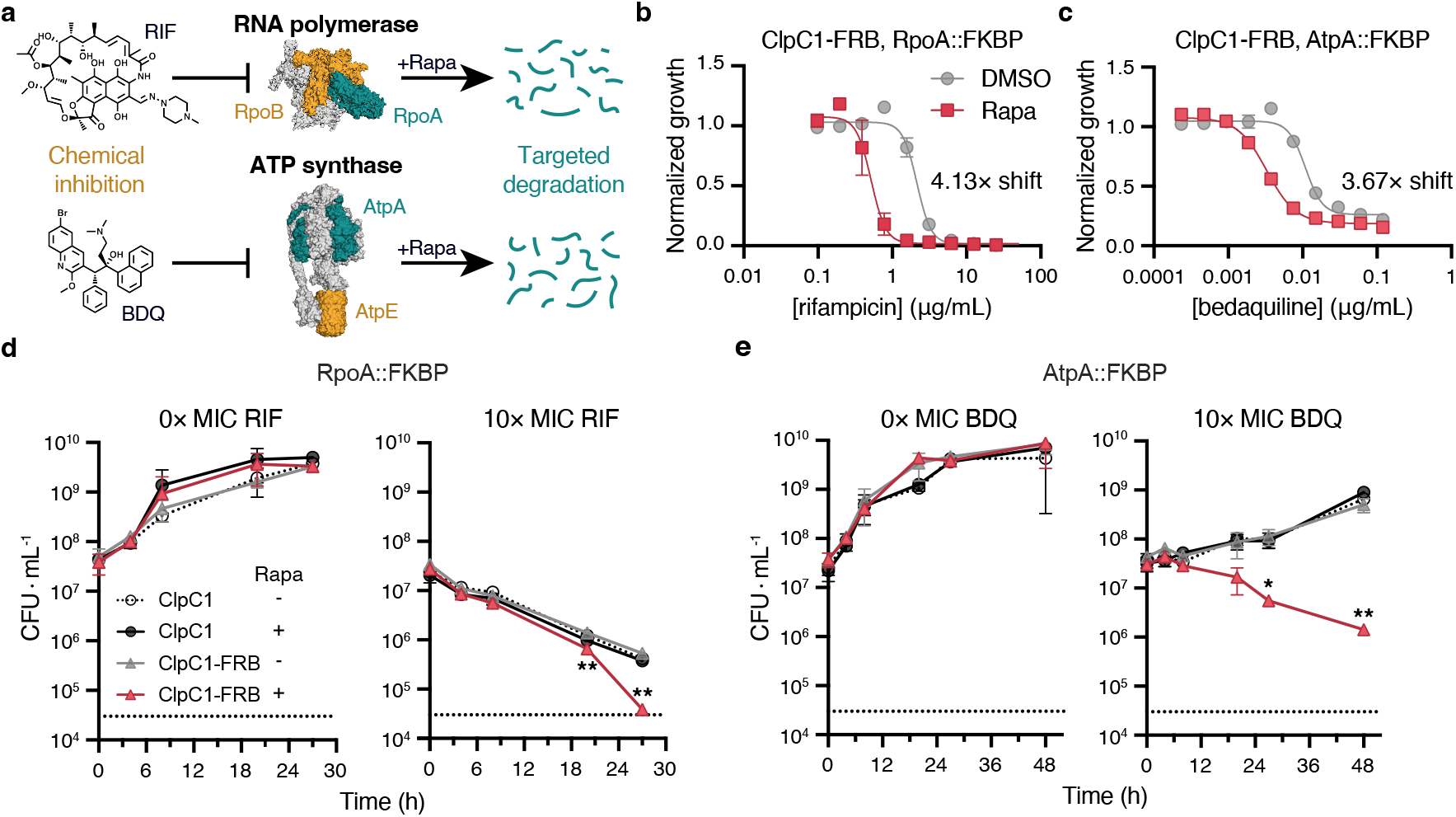
Targeted degradation of complex components sensitizes to and enhances killing by antibiotics targeting the same complex. **a** Schematic of combination approach to potentiate antibiotic efficacy, in which different members of the multi-component complexes RNA polymerase (top, PDB: 5UHA) or ATP synthase (bottom, PDB: 7NJK) are both inhibited with small molecule antibiotics (yellow) and targeted for degradation by rapamycin (green). **b-c** Half-maximal minimum inhibitory concentration (MIC_50_) dose response measuring the sensitivity of strains expressing RpoA-FKBP-FLAG (**b**) or AtpA-FKBP-FLAG (**c**) to the indicated antibiotics in media supplemented with DMSO or 0.5 μg ml^-1^ rapamycin. **d**-**e** Kill curves measuring the number of individual colony forming units (CFU) following treatment of strains expressing RpoA (**d**) or AtpA (**e**) with 0× MIC (left) or 10× MIC (right) the corresponding antibiotic for the indicated times. This assay measures the killing kinetics of antibiotics when supplemented with DMSO or 0.5 μg ml^-1^ rapamycin. *P* values were determined by unpaired two-tailed *t*-tests and compared 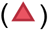 with 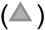. \**P* < 0.05, \*\**P* < 0.01. Exact *P*-values: (**e**) t = 20 h, \*\**P* = 0.0049; t = 27 h, \*\**P* = 0.0017, (**f**) t = 27 h, \**P* = 0.0101; t = 48 h, \*\**P* = 0.0055. For **b-e**, data are mean ± s.d. of three technical replicates and are representative of two independent experiments.

In addition to inhibition of growth (i.e., bacteriostatic activity), we further asked whether targeted degradation of RpoA and AtpA specifically enhances bactericidal killing of cells by rifampicin and bedaquiline, respectively. While degrading RpoA had no observable effect on bacterial count in the absence of rifampicin, when treated with 10× the observed MIC_50_ of rifampicin, targeted degradation of RpoA resulted in significantly greater cell killing compared to cells with no degradation (Fig. 5d, *P* < 0.01 for 20 and 27 hours). Similarly, degrading AtpA had no impact on bacterial count without bedaquiline, but treatment with 10× the MIC_50_ of bedaquiline resulted in significantly enhanced cell killing compared to cells with no degradation, which did not demonstrate any killing at all (Fig. 5e, *P* < 0.05 for 27 and 48 hours). This synergy is particularly striking as bedaquiline has been shown to have delayed bactericidal activity clinically^29,30^. Taken together, our findings highlight current antibiotic targets as pathways which can also be attacked by ClpC1-mediated targeted protein degradation, yielding direct effects on reducing bacterial growth, but importantly also sensitizing bacteria to currently approved antibiotics and accelerating their killing kinetics.

## Discussion

In this work, we developed and validated a platform for the systematic identification of promising targets in the development of BacPROTAC targeted degrader antibiotics against mycobacteria. Genetic fusions to the FRB and FKBP domains enable the induced proximity of targets and the ClpC1 ATPase with the small molecule rapamycin, facilitating the subsequent proteolytic degradation of endogenous proteins. We show that most but not all proteins we screened as ClpC1 targets are effectively degraded using this approach, a subset of which demonstrate growth inhibitory, as well as antibiotic sensitizing and killing rate enhancement effects on *Msm*. Our results demonstrate the potential of TPD as a new modality for antimicrobial therapeutics.

TPD offers some advantages over traditional antibiotics. Since active site inhibition is not necessary for this modality, molecules which may be poor inhibitors could be repurposed as high affinity ligands for PROTAC development. Additionally, not being limited to an enzymatic active site expands the repertoire of 1) potential targets to include those “undruggable” by traditional means, as well as that of 2) potential ligands that can be identified for a given target (that is, while an enzyme has only one active site, high affinity binders can be raised to multiple sites on a protein). The latter point suggests that even when resistant mutants eventually arise to a TPD antibiotic, a different ligand could be swapped that bound the target protein at a different site. Moreover, the bifunctional nature of targeted degraders provides an alternative means of specificity.

Though we ultimately selected the ClpC1P1P2 proteolytic complex as the mediator of TPD in mycobacteria in our system, there are other endogenous protein degradation systems that may also be suitable. The Pup proteasome system (PPS) is more analogous to PROTACs, as the PPS uses the pupylating ligase PafA which could reduce the necessary residence time of the molecule to achieve an effect, as necessary residence time is likely lower for pupylation at a single exposed histidine residue (PafA) compared to pulling in and unfolding full proteins (ClpC1). The PafA ligase and the proteasome likely have their own specificities, however, and like the ClpC1 targeting systems, might be appropriate for a restricted class of proteins.

When selecting targets for a TPD strategy, access to the target surface by the degradation mediating machinery (e.g., E3 ligase for PROTACs) is essential. The geometry of the nascent ternary complex formed when targets are brought in proximity to degradation machinery is therefore a critical factor determining whether proteins can be degraded or not^31^. In addition to the limitations on our ability to vary the geometry of potential targets, an additional layer of complexity to our ClpC1-mediated approach is that ClpC1 degradation has its own structural determinants. The rules governing whether ClpC1 will degrade a given protein are not well known, though previous work has observed that putative ClpC1 substrates tend to have relatively disordered termini^28^. Given differences in protein tertiary structure and termini composition among different proteins, it is not overly surprising that we observed a gradient of degradability among selected target proteins. While the proteins we selected span a diversity of essential functions and subcellular locations, the number of proteins we screened (n=17) is too small to make any meaningful associations between protein features and degradability per se; further studies that screen the full set of essential mycobacterial genes would be able to answer this question.

Interestingly, while we had expected to find that being a predicted native ClpC1 substrate^28^ would correlate with effective TPD in our system, we found that the predicted substrates were broadly distributed across the degradability spectrum, with the two most intractable mycobacterial proteins, EttA and InhA, both being predicted to be native substrates. Meanwhile, that we were able to degrade 6/6 proteins not predicted to be native ClpC1 substrates perhaps suggests that ClpC1 has some promiscuity in its ability to degrade proteins delivered to it, further highlighting this proteolytic complex component as a promising mediator of TPD in mycobacteria. This suggests that the inherent degradability of a given target by induced proximity is likely due to the confluence of multiple factors.

By microscopy, we noted that for some, but not all targets, rapamycin addition resulted in the formation of fluorescent puncta. This raises the possibility that delivery of target proteins to ClpC1 could have a dual effect: first, proximity-mediated degradation of that protein, and second, forced removal of that protein from its native localization, potentially perturbing optimal function or complex formation. This forced removal may be sufficient to cause unfavorable outcomes in bacteria even without substantial degradation; for example, Ffh was only degraded by about 30% of its steady state levels, yet we observed substantial growth delays on solid media and significantly reduced colony formation when degraded. There is some evidence for this concept in *Mtb*, where it has been shown that small molecules that disrupt a native protein-protein interaction shows strongly bactericidal activity^32^.

In our study, were we able not only to show effective protein degradation using our system but also to demonstrate TPD-dependent growth inhibition for some targets; furthermore, we show that TPD can mediate antibiotic sensitization and more rapid antibiotic killing for other targets. The strongest of these synergistic phenotypes was with AtpA depletion enhancing the rate of killing by bedaquiline, one of the newest antibiotics for the treatment of multi-drug resistant *Mtb* infection^33,34^. Bedaquiline has been previously shown to have delayed killing kinetics, which are not well understood and finding avenues to shorten this period – as we showed with AtpA degradation – could prove advantageous in a clinical setting. The phenotypes we observed with AtpA degradation was a curious example due to the differential growth phenotypes we observed for different members of the ATP synthase complex. Though targeted degradation of AtpA results in reduced colony formation with substantial growth delays on solid media, no such reduction in colony formation or growth delay is observed for AtpG, AtpD, or AtpH degradation. This would suggest that either 1) we achieve differential levels of protein degradation among the different subunits, or 2) mycobacteria are particularly susceptible to AtpA depletion. Recent work that systematically characterized the effect of essential genes’ transcript depletion by CRISPRi on bacterial fitness in *Mtb* and *Msm^35^* found that *Msm* is similarly “vulnerable” to depletion of all four ATP synthase subunits, suggesting that in the strains with *fkbp* fusions at their chromosomal loci, we are able to target a higher percentage of cellular AtpA for degradation compared to the other subunits. Our study demonstrates that the removal of a highly “degradable” target is not inherently sufficient to result in growth inhibitory (bacteriostatic) or cell lethality (bactericidal) phenotypes, despite these proteins being identified as essential^11^. This could be because the new steady state level of protein after TPD is still sufficient to perform cellular function under our culture conditions. The availability of the recent CRISPRi vulnerability dataset now provides a resource which can guide the selection of additional TPD targets, enriching for proteins to which cells are particularly susceptible when even small amounts are depleted.

An important limitation of this study is that we were only able to fuse ClpC1 with FRB at its C-terminus. Therefore, delivery of target proteins by rapamycin addition is not to the hexameric pore, but rather the base of the hexamer. We reason that what degradation we observed was due to the inherent on/off kinetics of the FRB-rapamycin-FKBP interaction increasing the local concentration of the target proteins around ClpC1, driving enhanced degradation. When employing a small molecule ligand that binds to the ClpC1 N-terminal domain (NTD), we expect greater degradation efficiency. Notably, the ClpC1-targeting ligands in the BacPROTAC work^10^ directly target the ClpC1 NTD; therefore, we expect that the potent targets identified in our work may be even more efficiently degraded with a cognate BacPROTAC molecule. Another limitation is that the growth and viability phenotypes following TPD we report have been generated in *Msm* cells, using genetic fusions to the FRB and FKBP domains (i.e., not a heterobifunctional small molecule). To enable further use of this system, it will be important to identify 1) small molecule or chemical fragment ligands to the *Mtb* orthologs of promising targets we identified and 2) ligands that bind optimally to *Mtb*ClpC1, enabling the generation of BacPROTACs to direct the degradation of native *Mtb* proteins.

Together, our work reports a platform that enables the rapid screening of bacterial target proteins for TPD and the identification of particularly labile targets that elicit antibacterial phenotypes. Further screening of both new and old therapeutic targets for suitability as TPD targets could be used for the development of new degrader BacPROTAC antibiotics. In particular, the demonstrated sensitization to and enhanced killing by existing antibiotics could mean that, when coupled with existing antibiotics, combination therapy with a BacPROTAC antibiotic might shorten *Mtb’s* requisite lengthy treatment times or even restore antibiotic sensitivity to drug-resistant strains.

## Methods

### Bacterial strains and culture conditions

*Mycobacterium smegmatis* mc^2^155 and derivative strains were grown in Middlebrook 7H9 liquid media (Millipore) supplemented with ADC (5 g l^-1^ bovine albumin fraction V, 2 g l^-1^ dextrose, 0.003 g l^-1^ catalase, 0.85 g l^-1^ NaCl), 0.2% (v/v) glycerol, and 0.05% (v/v) Tween-80 or plated on LB agar or 7H10 agar. Antibiotic selection concentrations for *Msm* were: 25 μg ml^-1^ kanamycin, 20 μg ml^-1^ nourseothricin, 50 μg ml^-1^ hygromycin, and 20 μg ml^-1^ zeocin. Antibiotic selection concentrations for *Escherichia coli* were: 50 μg ml^-1^ kanamycin, 40 μg ml^-1^ nourseothricin, 100 μg ml^-1^ hygromycin, and 50 μg ml^-1^ zeocin. All strains were cultured at 37°C. Rapamycin (Sigma) was dissolved in DMSO to a stock concentration of 10 mg ml^-1^.

### Strain construction

#### mc^2^155 Δ*clpC1* L5::*clpC1-frb* strain

The *clpC1-frb* strain was made by L5 allele swap. First, a *clpC1* (MSMEG_6091) merodiploid strain was generated by amplifying *clpC1* with 500 bp of the sequence upstream of the gene to capture the native promoter, assembling into a kan^r^ plasmid with the L5 integrase, and integrating as a single copy at the L5 phage chromosomal attachment site (*attB*)^36^. Native *clpC1* was knocked out by mycobacterial recombineering (phage-mediated homologous recombination)^37^, replacing the native *clpC1* allele with a zeocin selection marker, flanked by *loxP* sites. The knockout construct comprised 500 bp homology arms upstream and downstream of *clpC1* surrounding the floxed zeocin selection marker and was assembled into a pL5 vector^38^. Next, *clpC1* (with native promoter) and *frb* codon-optimized for mycobacteria were amplified, assembled into a ntc^r^ plasmid with the L5 integrase, and transformed into the *ΔclpC1 L5::clpC1* strain; successful swaps acquired nourseothricin resistance at the expense of kanamycin resistance, whereas double integrants contained both markers. A 1:1 ratio of single to double integrants indicates that there is no fitness advantage to either allele.

#### Fluorescent *target-fkbp-egfp* fusions

The genes listed in Fig. 3d, as well as *fkbp* and *egfp* codon optimized for mycobacteria were amplified, assembled under control of the high-strength P_UV15_ or P_smyc_ promoters (except for HupB and Wag31; medium-strength Pimyc promoter) into a kan^r^ plasmid with the Tweety integrase, and transformed in the WT *clpC1* or *ΔclpC1 L5::clpC1-frb* background.

#### Chromosomal locus *target-fkbp* fusions

The genes listed in Fig. 4d were tagged with *fkbp-flag* codon optimized for mycobacteria at their native locus by recombineering, by adding the coding sequence in frame immediately before each gene’s stop codon. The knockin construct comprised 500 bp homology arms upstream and downstream of each gene’s stop codon surrounding the floxed zeocin marker and was assembled into a pET-26b+ vector. The zeocin marker was removed by transformation with a Cre plasmid that is sucrose curable by SacB^39^. All chromosomal locus tags were with *fkbp-flag*, except *ffh*, which was *fkbp-egfp*. All plasmids used in this work were cloned using isothermal assembly^40^ and plasmid maps are available upon request.

#### Transformation

*E. coli* competent cells were prepared using rubidium chloride and transformed by heat shock. Electrocompetent *Msm* cells were prepared by growing cells to mid-log phase (OD_600_ ≈ 0.5), washing 3× with 10% (v/v) glycerol pre-chilled to 4°C, then resuspending in 1/100^th^ of the initial culture volume with 10% glycerol. DNA (200 ng for integrating plasmids, 1 μg for recombineering) was added to 100 μl competent cells and incubated on ice for 5 minutes, followed by electroporation in a 2 mm cuvette at 2500 V, 125 Ω, 25 μF using an ECM 630 electroporator (BTX). 7H9 media (1 ml) was added to the cells, which recovered for 3 h with shaking at 37°C, before spreading on LB agar and incubating at 37°C for 2-3 days.

### NanoLuc assay

To validate rapamycin-mediated FRB-FKBP dimerization in mycobacteria, *Msm* cells were grown to mid-log phase (OD_600_ ≈ 0.5) and plated at a final OD_600_ = 0.1 with 5-fold serial dilutions of rapamycin. The Nano-Glo^®^ 5× reagent (furimazine substrate + buffer; Promega) was prepared and added to each sample. Luminescence was measured in opaque white plates after incubation with linear shaking at 37°C for 5 min using a Spark 10M plate reader (Tecan).

### Protein structure modeling

The crystal structure of the stabilized mutant *Mtb*ClpC1 hexamer (PDB: 8A8U)^20^ was modeled using PyMOL (Schrodinger). The *Mtb*ClpC1 NTD trimer was modeled by aligning three individual structures of the *Mtb*ClpC1 NTD (PDB: 6PBS)^41^ with a *E. coli* ClpB NTD trimer (PDB: 6OG3)^42^ as previously in ref. 20.

### Protein expression and purification

The *Mtb clpC1* (Rv3596c) allele and *frb* codon optimized for mycobacteria were amplified and assembled to encode both N- and C-terminally fused gene products under control of the T7 promoter into a pET-26b+ plasmid, which encodes a 6xHis tag at the C-terminal end of the gene product. Constructs were transformed into BL21-CodonPlus(DE3)-RP cells (Agilent), which enables efficient expression of GC-rich genes in *E. coli*.

BL21 strains were grown in LB broth (Lennox) with shaking at 37°C, until they reached OD_600_ ≈ 0.8; protein expression was induced overnight with 100 μM IPTG and incubating with shaking at 18°C overnight. Cells were harvested by centrifugation, subjected to a −80°C freeze-thaw cycle, and resuspended in protein binding buffer (25 mM Gomori buffer pH 7.6, 100 mM KCl, 5% (v/v) glycerol). Next, cells were treated with 12.5 μg ml^-1^ lysozyme for 5 min and then lysed by sonication (30% power pulses for 30 s, alternating with ice incubation for 1 min; repeat 4×). Cell debris was removed by centrifugation and His-tagged ClpC1 proteins were purified from supernatants with Ni-NTA agarose beads (Qiagen). After washing in binding buffer supplemented with 20 mM imidazole, proteins were eluted with binding buffer supplemented with 100 mM and 200 mM imidazole. Input, flow through, and both elutions were examined by SDS-PAGE by loading samples in a NuPAGE 4-12% Bis-Tris precast gradient gel (Invitrogen) using MES SDS buffer.

Eluted proteins were concentrated and desalted with an Amicon^®^ Ultra-4 Centrifugal Filter Device (30 kDa MWCO, Millipore Sigma), with equilibration in protein binding buffer lacking imidazole; protein concentrations were ascertained with the Pierce™ Coomassie Plus (Bradford) Assay Reagent (ThermoScientific) and comparison to a BSA standard curve. Purified fractions were pooled, aliquoted, and stored at −80°C. FRB-tagged *Mtb*ClpC1 purification was conducted as one independent experiment and WT *Mtb*ClpC1, *Mtb*ClpP1, and *Mtb*ClpP2 were purified previously^21^.

### *In vitro* assays

#### ClpC1P1P2 protease activity

Degradation of the eGFP-ssrA by purified *Mtb*ClpC1P1P2 was measured by examining loss in fluorescence signal over time. *Mtb*ClpC1 (WT and FRB-tagged proteins, 0.26 μM hexamer), *Mtb*ClpP1P2 (0.26 μM tetradecamer), and eGFP-ssrA (1.25 μM) were mixed in buffer containing: 25 mM Gomori buffer pH 7.6 supplemented with 100 mM KCl (Sigma), 5% glycerol (Sigma), 8 mM Mg-ATP (Sigma), and 2.5 mM Bz-Leu-Leu^43^ (Laboratory of Dr. William Bachovchin, Tufts University). Fluorescence signal decay was measured (Ex/Em: 485 nm / 520 nm) continously in a SpectraMax M5 microplate reader (Molecular Devices) at 37°C for 35 min.

#### ClpC1 ATPase activity

ATP hydrolysis by purified *Mtb*ClpC1 was measured using a pyruvate kinase and lactate dehydrogenase (PK/LDH) coupled enzymatic assay in which absorbance at 340 nm is an inverse proxy for the rate of ATP hydrolysis. *Mtb*ClpC1 (WT and FRB-tagged proteins, 0 − 0.7 μM hexamer), 1 mM phosphoenolpyruvate (Sigma), and 20 U ml^-1^ PK/LDH (Sigma) were mixed in buffer containing: 50 mM Tris pH 7.6 (Sigma), 100 mM KCl (Sigma), 1 mM DTT (Sigma), 1 mM NADH (Sigma), and 4 mM Mg-ATP (Sigma). Absorbance_340_ was measured continuously in a SpectraMax M5 microplate reader (Molecular Devices) at 37°C for 32 min.

### Flow cytometry

To measure the fluorescence of the target reporter fusions in the presence of rapamycin, *Msm* cells were grown to mid-log phase (OD_600_ ≈ 0.5) and plated at a final OD_600_ = 0.2 (for the time course flow cytometry, OD_600_ = 0.1) with the denoted concentration of rapamycin or DMSO (vehicle control). Cells were incubated with shaking at 700 rpm at 37°C for the specified times (0 h, 3 h, 6 h, 9 h, 24 h), diluted in 7H9 media (to remain under 10,000 events sec^-1^), and fluorescence was quantified on a MACSQuant^®^ VYB flow cytometer using the green fluorescence laser and filter set (channel B1, Ex/Em: 488 nm / 525/50 nm) with at least 20,000 events captured for each sample. To suppress spurious events due to noise or cell debris, event triggers were applied on forward scatter peak height (FSC-H > 0.8) and side scatter area (SSC-A > 1.2). To exclude cellular aggregates and morphological outliers, gates were applied with FlowJo (v. 10.8), and a representative gating strategy can be found in Supplementary Figure 6a. The remaining events were normalized to mode and presented as histograms displaying log10 transformed green fluorescence intensity area (AlexaFluor488-A), using GFP signal as a proxy for target protein levels *in vivo*. Full flow cytometry plots for all 17 targets and time points summarized in Fig. 3b can be found in Supplementary Figure 7a. For the time course flow cytometry, cells were prepared and then replica plated for each time point. At least 20,000 events were captured; after gating for single *Msm* cells, at least 15,000 cells remained for each sample measured. Due to plate size limitations, technical replicates were measured from the same sample.

### Immunoblotting

To directly examine protein levels when redirected to ClpC1-FRB, *Msm* cells were grown to mid-log phase (OD_600_ ≈ 0.5), pelleted by centrifugation, and resuspended in 1× TBS supplemented with the cOmplete™ Protease Inhibitor Cocktail (Roche). Resuspended cells were aliquoted into FastPrep^®^ 2 ml Lysing Matrix tubes (MPBio) and lysed by beat beating with a Precellys 24 tissue homogenizer (Bertin Technologies; 6,500 rpm for 45 s, alternating with ice incubation for 2 min; repeat 3×). Cell debris was removed by centrifugation, supernatant protein content was evaluated by NanoDrop 1000 (ThermoScientific), and all samples were normalized to the same protein concentration in 1× TBS with protease inhibitor. Normalized protein aliquots were treated with TURBO™ DNase (Invitrogen) for 15 min at 37°C to digest mycobacterial genomic DNA and then denatured with boiling in Laemmli buffer. For all samples, 140 μg of protein was loaded onto NuPAGE 4-12% Bis-Tris precast gradient gels (Invitrogen) using MES SDS buffer, transferred to PVDF membranes using a Trans-Blot Turbo Transfer System (Bio-Rad), and blocked with 1:1 SEA-BLOCK blocking buffer (ThermoScientific) and 1 × PBS with shaking for 1 h at 25°C. Membranes were probed with the specified antibodies in 1:1 SEA-BLOCK and 1 × PBS-Tween (phosphate-buffered saline with 0.1% (v/v) Tween-20) with shaking for 1 h each primary and secondary at 25°C. Immunoblots were imaged using the Odyssey^®^ CLx Infrared Imaging System (LI-COR) and processed using LI-COR ImageStudio. The full uncropped scans of the blot shown in Fig. 2E can be found in Supplementary Figure 5. The mouse IgG2b monoclonal RpoB antibody (Invitrogen, #8RB13) was diluted 1:1,000. The rabbit IgG monoclonal GFP antibody (Abcam, #EPR14104) was diluted 1:1,000. The IRDye^®^ 680RD goat IgG (H+L) anti-mouse and IRDye^®^ 800CW goat IgG (H+L) anti-rabbit (LI-COR) fluorescent secondary antibodies were diluted 1:15,000.

### Time-lapse fluorescence microscopy and data analysis

To generate the time-lapse data for targeted degradation of RpoA-FKBP-eGFP in Fig. 2f, cells were grown until late log phase (OD_600_ ≈ 1.0) and seeded on 2.0% agarose pads supplemented with 7H9. The agarose pad was cast in a customized plastic frame and placed in a low-evaporation imaging disk as described previously^27^. Fluorescence and phase-contrast images were acquired every 10 min over a 9 h period with a Plan Apo 60 3 1.45 NA objective using a Nikon Ti-E inverted, widefield microscope equipped with a Nikon Perfect Focus system with a Piezo Z drive motor, Andor Zyla sCMOS camera, and an Agilent MLC400 Monolithic laser combiner. Fluorescence signals were acquired using a 6-channel Spectra X LED light source and the Sedat Quad filter set. The excitation and emission filters used for the green fluorophores (eGFP) were Ex. 470/24 nm and Em. 515/25 nm. Time-lapse snapshots were rendered using customized Python scripts. To analyze time-lapse data, our previously established microscopy image analysis pipeline MOMIA^27^ was modified to track the fluorescence of individual cells and emergence of new cell generations.

### Target protein microscopy image acquisition

To capture the microscopy data for targeted degradation of all 17 target proteins examined in Fig. 3f and Supplementary Figure 8a, cells were grown to mid-log phase (OD_600_ ≈ 0.5) and plated at a final OD_600_ = 0.1 and incubated with the indicated concentration of rapamycin for 9 h with shaking at 700 rpm at 37°C. Cells were seeded on 2.0% agarose pads; fluorescence and phase-contrast images were acquired using the same microscope configurations as used for the time-lapse experiments. Image crops were rendered using customized Python scripts and the dynamic range used for each individual target is reported in Supplementary Figure 8b.

### Quantitative imaging of colony growth

To track colony growth dynamics, cells were grown to early-log phase (OD_600_ ≈ 0.3), then serially diluted and plated on solid 7H10 media supplemented with ADC and 0.5 μg ml^-1^ rapamycin or the equivalent volume of DMSO. Plates were incubated for 2-3 days at 37°C and images were captured with a 12-megapixel camera with a 120° FOV at 0, 6, 8, 13, 21,25, 30, 36, 48, and 56 h after plating. Our previously devised colony tracing pipeline^44^ was augmented with two deep learning models^45^ that were trained to automate the identification of the grid lines of each plate and also to perform semantic segmentation of plate photos. The pipeline was applied as previously to track the expansion of individual colony areas over time and was repurposed to generate CFU counts displayed in Fig. 4d and Supplementary Figure 11a.

### Minimum inhibitory concentration (MIC) assay

To calculate their dose response to antibiotics and targeted degradation, *Msm* cells were grown to mid-log phase (OD_600_ ≈ 0.5) and plated at a final OD_600_ = 0.0015 in each well of a 96-well plate containing serial dilutions of the specified antibiotics. For assays with concurrent targeted degradation, all wells were supplemented with 0.5 μg ml^-1^ rapamycin or DMSO. Plates were incubated with drug with shaking at 150 rpm at 37°C overnight. Resazurin was added to a final concentration of 0.002% (w/v), plates were incubated up to 18 h, and fluorescence conversion was measured (Ex/Em: 560 nm / 590 nm) in a SpectraMax M2 microplate reader (Molecular Devices). Background fluorescence was subtracted from wells without cells and fluorescence signal was normalized to wells without any compound. A nonlinear regression was used to fit a sigmoid curve to the dose response data and calculate the half-maximal minimum inhibitory concentration (MIC_50_, GraphPad Prism).

### Antibiotic kill kinetics curves

To examine the effect of targeted degradation on antibiotic killing, *Msm* cells were grown to midlog phase (OD_600_ ≈ 0.5) and plated at a final OD_600_ = 0.08 in 24 well plates; wells were supplemented with 0× or 10× their measured MIC of rifampicin or bedaquiline and with 0.5 μg ml^-1^ rapamycin or DMSO. Plates were incubated for 2 days with shaking at 155 rpm at 37°C and cells were sampled at 0, 4, 8, 20, 27 (both antibiotics), and 48 h (bedaquiline only) after plating. Sampled cells were serially diluted and plated on plain LB agar; plates were incubated for 2-3 days at 37°C and individual colony forming units (CFUs) were counted.

### Bacterial growth kinetics curves

To examine the effect of targeted degradation on bacterial growth in liquid media, *Msm* cells were grown to mid-log phase (OD_600_ ≈ 0.5) and plated at a final OD_600_ = 0.0015. Wells were supplemented with the specified concentration of rapamycin or DMSO. Absorbance (OD_600_) and/or eGFP fluorescence (Ex/Em: 488 nm / 510 nm) was measured in clear 96 well plates sealed with a Breathe-Easy^®^ membrane (Sigma) with linear shaking at 37°C for 48 h using a Spark 10M plate reader (Tecan).

### Statistical analysis

The statistical analyses and correlation plots described in this study were performed using the python package SciPy^46^ or GraphPad Prism. All experiments were performed at least twice (biological replicates), unless otherwise noted. Technical replicates were measured within the same experiment from distinct samples, unless otherwise noted. Means were compared using a two-tailed Student’s *t*-test. *P* < 0.05 was considered significant. For the microscopy experiments at least 100 cells were analyzed for each shown image and the image data shown are representative of multiple fields. No statistical methods were used to determine sample sizes used in this work and the investigators were not blinded to sample identity.

## Data availability

The data that support the findings of this study are available from the corresponding author upon reasonable request.

## Code availability

All code used to generate data in this work are available at a public GitHub depository (https://github.com/jzrolling/ClpC1_TPD).

## Acknowledgments

We thank members of the Rubin and Sarah Fortune labs for helpful discussions during this work and Douaa Mugahid for thoughtful feedback on the manuscript. We thank Kenan Murphy for sharing the ClpC1-eGFP *Msm* strain. Funding: This material is based upon work supported by the NSF Graduate Research Fellowship under Grant No. DGE1745303 (HW), the Harvard GSAS Herchel Smith Graduate Fellowship (HW), the Marcus Urann Graduate Fellowship (HW), NIH/NIAID award U19 AI142735 (EJR), and funding provided by Grace Wang and Josef Tatelbaum.

## Author contributions

HW, JZ, and ER conceived and designed the study; HW, JZ, OK, and MW performed experiments; HW, JZ, and IW performed data analysis; OK, TA, MC, ER gave technical and conceptual advice; HW and JZ wrote the original draft; HW, JZ, MC, IW, OK, TA, MW, and ER reviewed and edited the manuscript; JZ and ER supervised this study.

## Correspondence

and requests for materials should be addressed to Eric Rubin.

## Supplementary Figure Legends

**Supplementary Figure 1 | Rapamycin does not restrict mycobacterial growth at the concentrations used in this work.** Half-maximal minimum inhibitory concentration (MIC_50_) dose response measuring the sensitivity of *<u>Msm</u>* strains to rapamycin. Primary concentration of rapamycin used in this work indicated by dotted line. Data are mean ± s.d. of three technical replicates and are representative of two independent experiments. Related to Fig. 1.

**Supplementary Figure 2 | C-terminally tagged *Mtb*ClpC1-FRB retains ATPase activity *in vitro*. a** SDS-PAGE of purified tagged *Mtb*ClpC1 proteins expressed in BL21 cells. Arrow denotes expected size of fusion proteins. I = input; FT = flow-through; E1 = elution 1, 100 mM imidazole; E2 = 200 mM imidazole. **b** Schematic of ATP/NADH coupled *in vitro* assay to measure ClpC1 ATPase activity. **c** Absorbance at 340nm measuring *in vitro* ATPase activity of WT ClpC1 (left), ClpC1-FRB (middle), or FRB-ClpC1 (left) with the indicated concentrations of each protein. For **c**, data are individually plotted measurements, normalized to time = 0 h, and are representative of two independent experiments. **b** Created with BioRender.com. Related to Fig. 1.

**Supplementary Figure 3 | The ClpC1-FRB strain is not growth-impaired compared to WT *Msm*.** Optical density of bacterial cultures at 600nm measuring the growth kinetics of the indicated *Msm* strains over time when supplemented with DMSO (left), 0.1 μg ml^-1^ rapamycin (middle), or 0.5 μg ml^-1^ rapamycin (left) with shaking at 37°C. Data are mean ± s.d. of three technical replicates and are representative of two independent experiments. Related to Fig. 1.

**Supplementary Figure 4 | Rapamycin re-localizes FKBP-eGFP. a** Live cell, wide-field fluorescence microscopy images of cells expressing ClpC1 tagged at its chromosomal locus with eGFP and treated with DMSO (top) or 0.1 μg ml^-1^ rapamycin (bottom). Scale bar, 5 μm. **b** Western blot analysis of ClpC1-eGFP. **c** Live cell, wide-field fluorescence microscopy images of cells expressing FKBP-eGFP in the WT *clpC1* (left) or *clpC1-frb* (right) background and treated with DMSO (top) or 0.1 μg ml^-1^ rapamycin (bottom). **a,c** Data are representative images selected from among 4 fields for each and are representative of two independent experiments. Normalized axial intensity of FITC signal across the cell widths of cells (N shown in each panel). Arrows highlight signal increases at the cell edges. Related to Fig. 2.

**Supplementary Figure 5 | Uncropped western blot from Fig. 2e.** Western blot analysis of RpoA-FKBP-eGFP with DMSO or 0.1 μg ml^-1^ rapamycin addition in the WT *clpC1* or *clpC1-frb* background. Related to Fig. 2.

**Supplementary Figure 6 | Flow gating strategy and RpoA degradation kinetics correlation plot. a** Representative flow gating strategy employed in this work. **b** Correlation plot comparing fluorescent signal loss kinetics of RpoA-FKBP-eGFP by time-lapse microscopy and flow cytometry. Time-lapse data represents the median fluorescent signal of all cells and flow data represents the mean fluorescent signal of two technical replicates; both are normalized to time = 0 h. Related to Fig. 3.

**Supplementary Figure 7 | Mycobacterial proteins are differentially degraded with varying kinetics. a** Fluorescence of live cells as a proxy for protein levels of indicated targets in the *clpC1-frb* background over time. Density matched log phase cells incubated with DMSO or 0.5 μg ml^-1^ rapamycin with shaking at 37°C for the indicated times. **b** Flow data from (**a**) transformed into normalized signal delay for all indicated targets following first order exponential decay kinetics. For **a**, data are two technical replicates, representative of two independent experiments, and normalized to the mode; **b**, data are individually plotted technical replicate measurements, representative of two independent experiments, and normalized to time = 0 h. Related to Fig. 3.

**Supplementary Figure 8 | Mycobacterial proteins are differentially degraded and sometimes re-localized with rapamycin. a** Live cell, wide-field fluorescence microscopy images of cells expressing selected target proteins with DMSO or 0.1 μg ml^-1^ rapamycin. Scale bar, 5 μm. **b** Dynamic range of GFP signal for each indicated strain. **c** Correlation plot comparing fluorescent signal intensity for each tested target by microscopy and flow cytometry.

Samples stratified by treatment with DMSO or 0.1 μg ml^-1^ rapamycin. For **a,** data are representative images selected from among 4 fields for each and are representative of two independent experiments. In **c,** Data are bounded by the 95% confidence interval; time-lapse data represents the median fluorescent signal of all cells and flow data represents the mean fluorescent signal of two technical replicates. Related to Fig. 3.

**Supplementary Figure 9 | Rapamycin stably directs degradation of RpsJ for 48 h. a-b** Cell density (OD_600_)-normalized fluorescence of live cells as a proxy for protein levels of FKBP-eGFP (**a**) or RpsJ-FKBP-eGFP (**b**) in the *clpC1-frb* background. Density matched log phase cells incubated with DMSO or 1 μg ml^-1^ rapamycin with shaking at 37°C. Data are individually plotted technical replicate measurements. Related to Fig. 3.

**Supplementary Figure 10 | Targeted degradation of RpoA and AtpA delays growth in liquid media. a-b** Optical density of bacterial cultures at 600nm measuring the growth kinetics of strains expressing RpoA (**a**) or AtpA (**b**) in the WT *clpC1* (left) or *clpC1-frb* (right) background when supplemented with DMSO or 10 μg ml^-1^ rapamycin with shaking at 37°C. Data are mean ± s.d. of three technical replicates and are representative of two independent experiments. Related to Fig. 4.

**Supplementary Figure 11 | Targeted degradation growth inhibition phenotypes are ClpC1-FRB dependent. a** Total colonies formed during outgrowth on solid media containing DMSO or 0.5 μg ml^-1^ rapamycin for the indicated targets in the *clpC1* background. Density matched log phase cells serially diluted, plated on solid media containing DMSO or 0.5 μg ml^-1^ rapamycin, and incubated at 37°C. *P* values were determined by unpaired two-tailed *t*-tests and compared 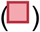 with 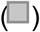. \**P* < 0.05, Exact *P*-value: SecA1, \*\**P* = 0.0214. **b** Quantitation of colony outgrowth dynamics by colony size tracking by area (mm^2^) of individual colonies of the indicated targets in the *clpC1* background over time. **c** Representative images illustrating colony outgrowth dynamics of the indicated targets in the *clpC1* background over time. Cells plated as in **a-b**. For **a**, data are mean ± s.d. of three technical replicates and are representative of two independent experiments; in **b**, dark lines are the mean of three technical replicates, are bounded by the 95% confidence interval, and are representative of two independent experiments. For **c**, images are representative of three technical triplicates and two independent experiments. Related to Fig. 4.

**Supplementary Figure 12 | *Msm* dose response to antibiotics with targeted protein degradation. a-d** Half-maximal minimum inhibitory concentration (MIC_50_) dose response measuring the sensitivity of indicated strains to rifampicin (**a**), bedaquiline (**b**), streptomycin (**c**), or linezolid (**d**) in media supplemented with DMSO or 0.5 μg ml^-1^ rapamycin. Observed fold-shifts are denoted on the corresponding plot. For **a-d**, data are mean ± s.d. of three technical replicates. Related to Fig. 5.

