## Supplementary Figures for "Chemically-induced targeted protein degradation in mycobacteria uncovers antibacterial effects and potentiates antibiotic efficacy"

#### Supplementary Figure 1

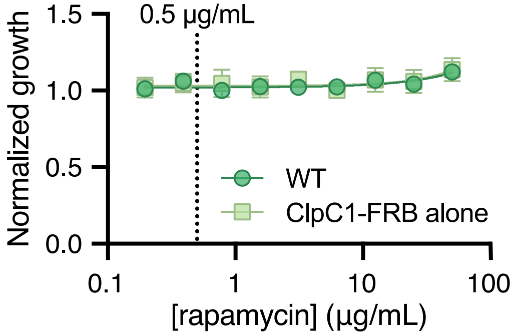


**Supplementary Figure 1 | Rapamycin does not restrict mycobacterial growth at the concentrations used in this work.** Half-maximal minimum inhibitory concentration ($MIC_{50}$) dose response measuring the sensitivity of *Msm* strains to rapamycin. Primary concentration of rapamycin used in this work indicated by dotted line. Data are mean ± s.d. of three technical replicates and are representative of two independent experiments. Related to Fig. 1.

### Supplementary Figure 2

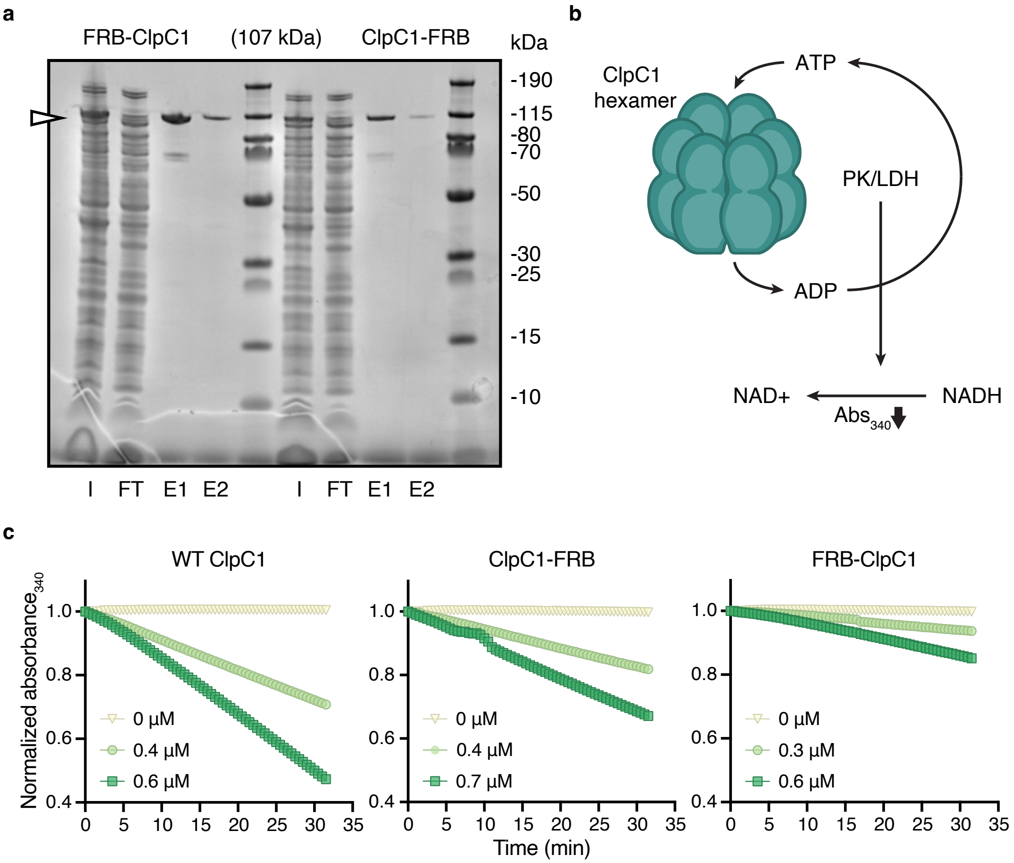


**Supplementary Figure 2 | C-terminally tagged *Mtb*ClpC1-FRB retains ATPase activity *in vitro.* a** SDS-PAGE of purified tagged *Mtb*ClpC1 proteins expressed in BL21 cells. Arrow denotes expected size of fusion proteins. I = input; FT = flow-through; E1 = elution 1, 100 mM imidazole; E2 = 200 mM imidazole. **b** Schematic of ATP/NADH coupled *in vitro* assay to measure ClpC1 ATPase activity. **c** Absorbance at 340nm measuring *in vitro* ATPase activity of WT ClpC1 (left), ClpC1-FRB (middle), or FRB-ClpC1 (left) with the indicated concentrations of each protein. For **c**, data are individually plotted measurements, normalized to time = 0 h, and are representative of two independent experiments. **b** Created with BioRender.com. Related to Fig. 1.

### Supplementary Figure 3

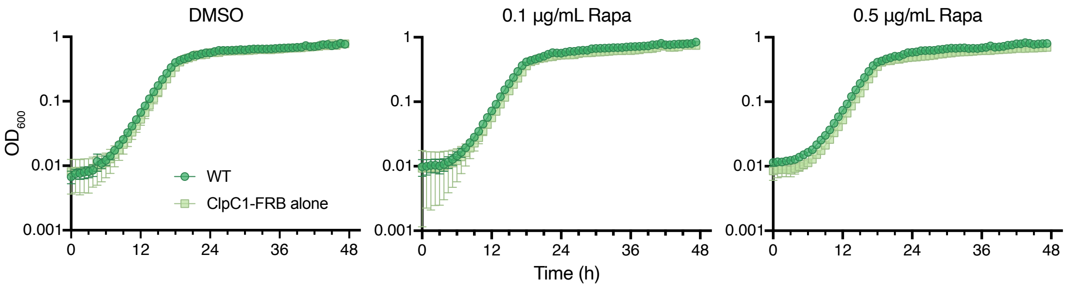


**Supplementary Figure 3 | The ClpC1-FRB strain is not growth-impaired compared to WT *Msm*.** Optical density of bacterial cultures at 600nm measuring the growth kinetics of the indicated *Msm* strains over time when supplemented with DMSO (left), 0.1 µg ml$^{-1}$ rapamycin (middle), or 0.5 µg ml$^{-1}$ rapamycin (left) with shaking at 37° C. Data are mean $\pm$ s.d. of three technical replicates and are representative of two independent experiments. Related to Fig. 1.

### Supplementary Figure 4

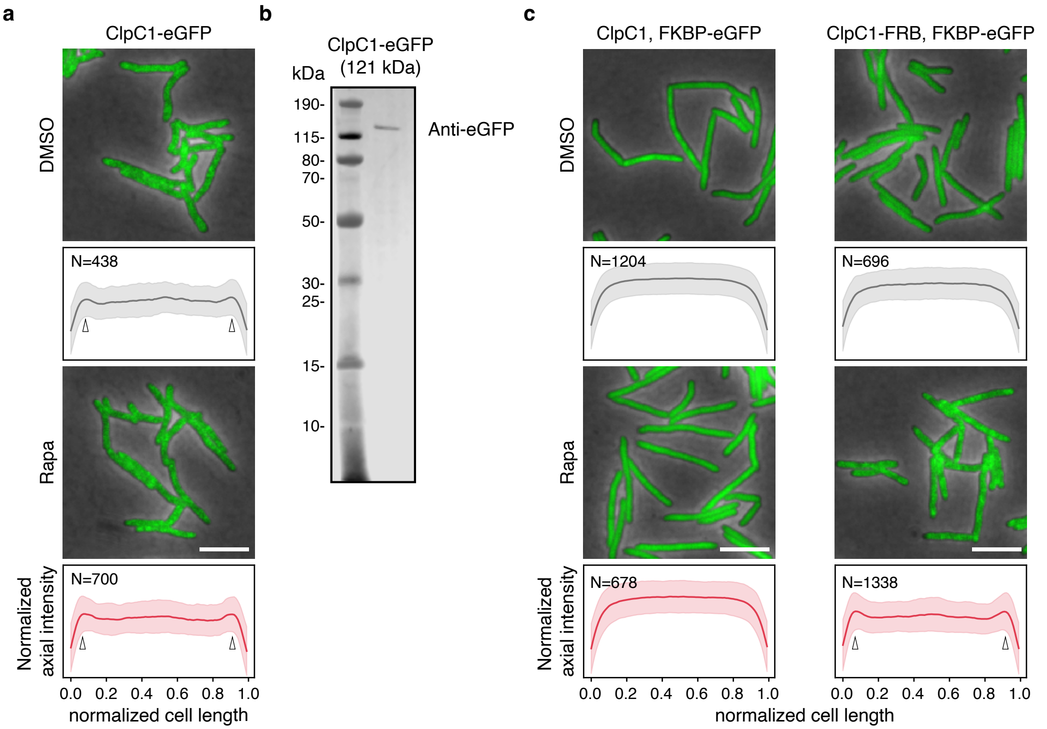


**Supplementary Figure 4 | Rapamycin re-localizes FKBP-eGFP. a** Live cell, wide-field fluorescence microscopy images of cells expressing ClpC1 tagged at its chromosomal locus with eGFP and treated with DMSO (top) or 0.1 μg $ml^{-1}$ rapamycin (bottom). Scale bar, 5 μm. **b** Western blot analysis of ClpC1-eGFP. **c** Live cell, wide-field fluorescence microscopy images of cells expressing FKBP-eGFP in the WT *clpC1* (left) or *clpC1-frb* (right) background and treated with DMSO (top) or 0.1 μg $ml^{-1}$ rapamycin (bottom). **a,c** Data are representative images selected from among 4 fields for each and are representative of two independent experiments. Normalized axial intensity of FITC signal across the cell widths of cells (N shown in each panel). Arrows highlight signal increases at the cell edges. Related to Fig. 2.

### Supplementary Figure 5

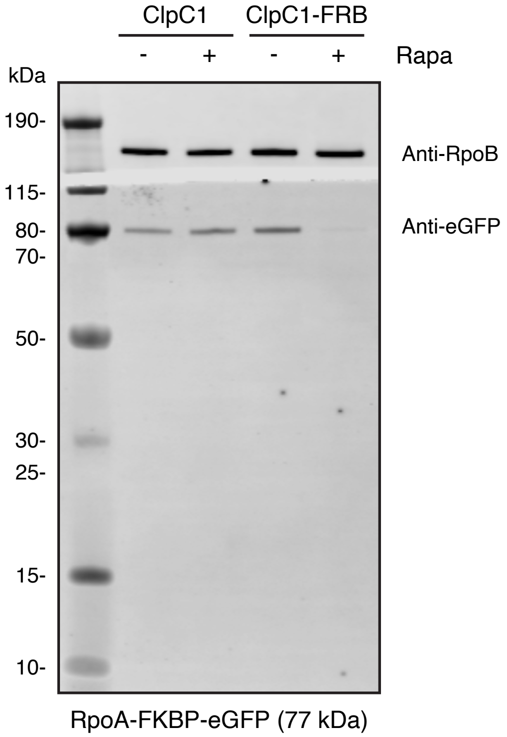


**Supplementary Figure 5 | Uncropped western blot from Fig. 2e.** Western blot analysis of RpoA-FKBP-eGFP with DMSO or 0.1 μg $ml^{-1}$ rapamycin addition in the WT *clpC1* or *clpC1-frb* background. Related to Fig. 2.

### Supplementary Figure 6

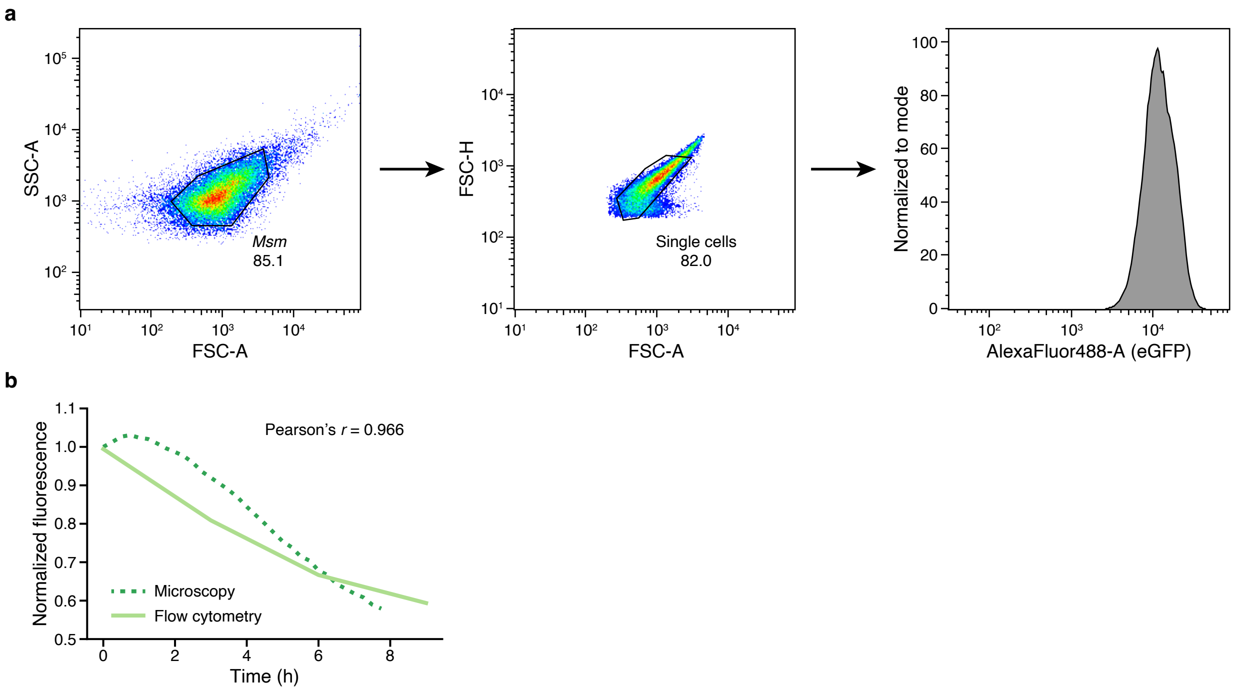


**Supplementary Figure 6 | Flow gating strategy and RpoA degradation kinetics correlation plot. a** Representative flow gating strategy employed in this work. **b** Correlation plot comparing fluorescent signal loss kinetics of RpoA-FKBP-eGFP by time-lapse microscopy and flow cytometry. Time-lapse data represents the median fluorescent signal of all cells and flow data represents the mean fluorescent signal of two technical replicates; both are normalized to time = 0 h. Related to Fig. 3.

### Supplementary Figure 7

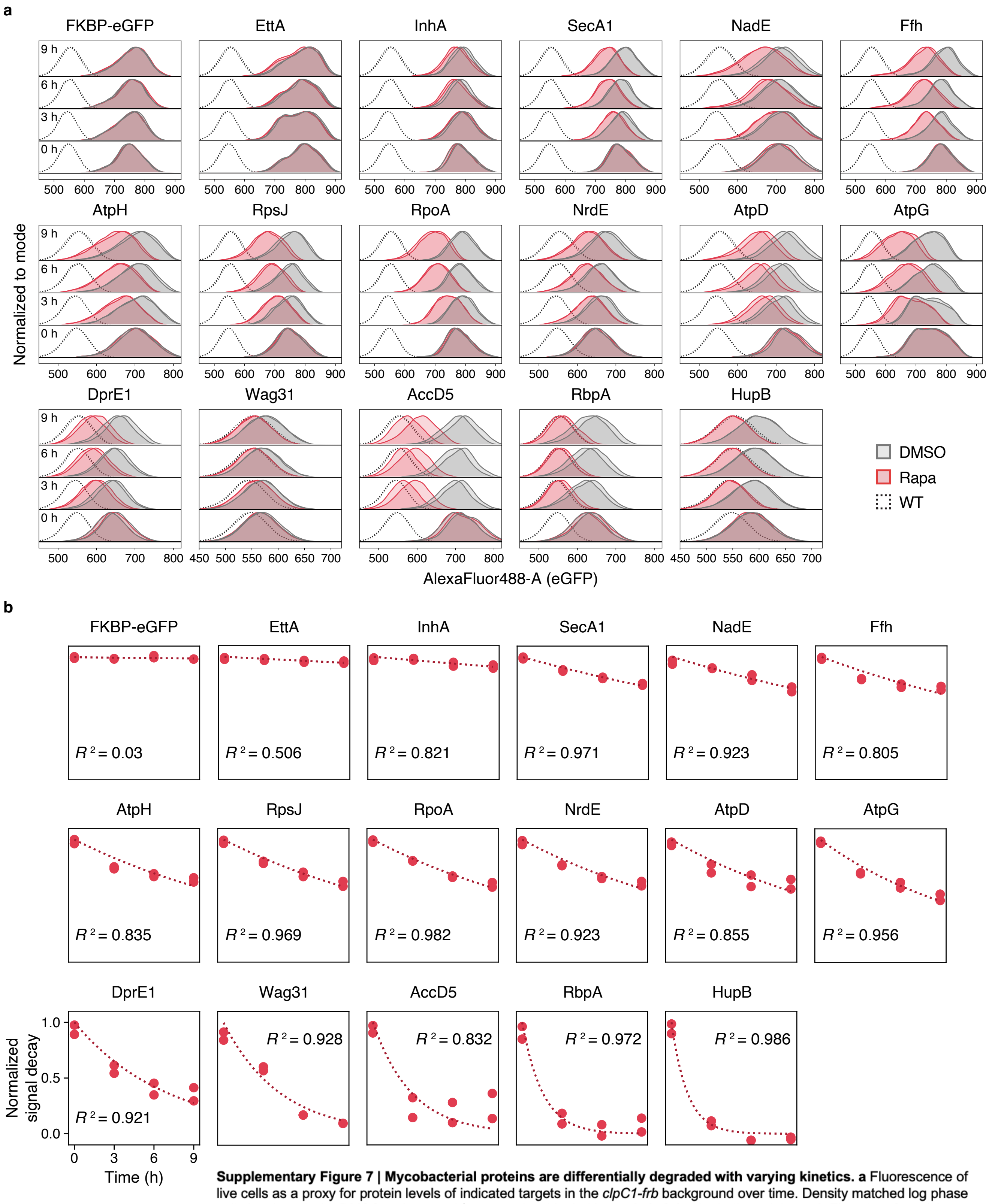


**Supplementary Figure 7 | Mycobacterial proteins are differentially degraded with varying kinetics. a** Fluorescence of live cells as a proxy for protein levels of indicated targets in the *clpC1-frb* background over time. Density matched log phase cells incubated with DMSO or 0.5 µg $ml^{-1}$ rapamycin with shaking at 37° C for the indicated times. **b** Flow data from (**a**) transformed into normalized signal delay for all indicated targets following first order exponential decay kinetics. For **a**, data are two technical replicates, representative of two independent experiments, and normalized to the mode; **b**, data are individually plotted technical replicate measurements, representative of two independent experiments, and normalized to time = 0 h. Related to Fig. 3.

#### Supplementary Figure 8

**a**

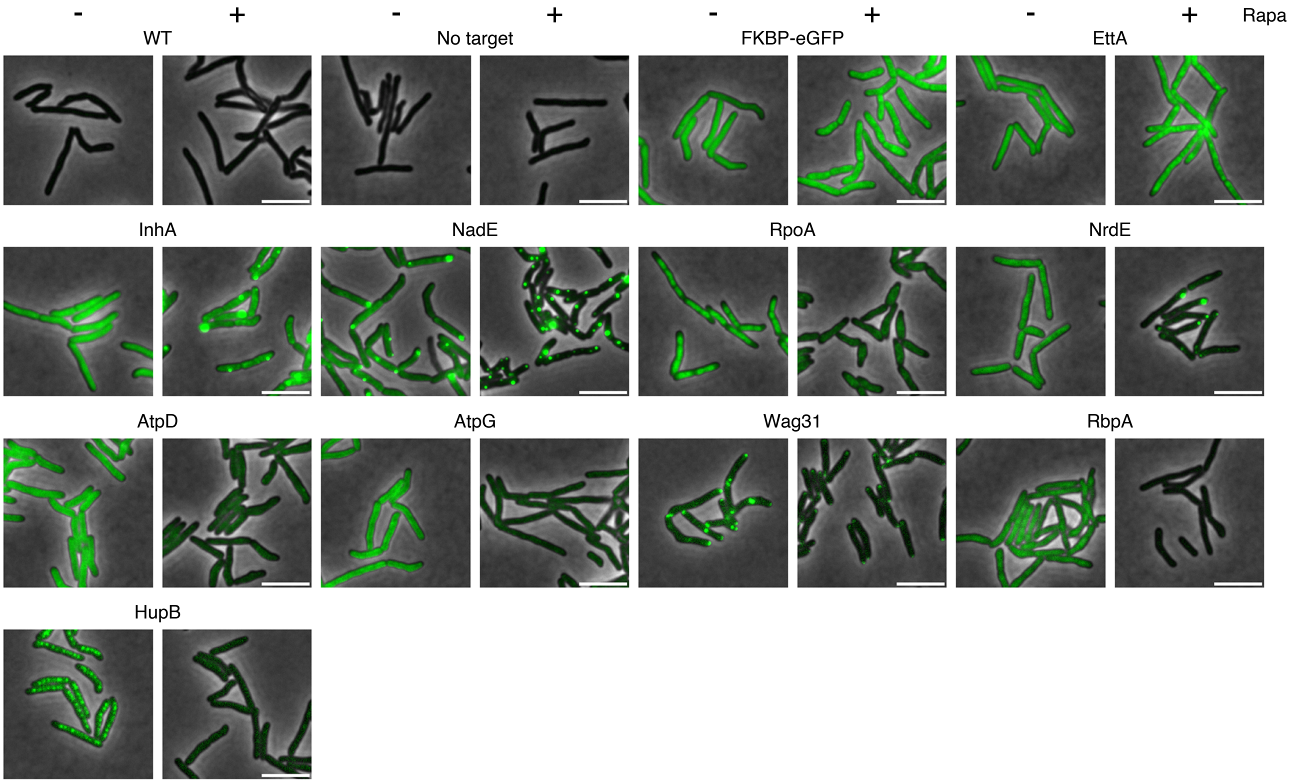


**b**

| Strain | dynamic range | Strain | dynamic range |
|---|---|---|---|
| WT | 500-1000 | RpoA | 500-4000 |
| ClpC1-FRB alone | 500-1000 | NrdE | 500-2500 |
| FKBP-eGFP | 500-4000 | AtpD | 500-2500 |
| EttA | 500-4000 | AtpG | 500-2500 |
| InhA | 500-5000 | DprE1 | 500-1000 |
| SecA1 | 500-5500 | Wag31 | 500-1000 |
| NadE | 500-4000 | AccD5 | 500-2000 |
| Ffh | 500-4000 | RbpA | 500-1000 |
| AtpH | 500-2500 | HupB | 500-1000 |
| RpsJ | 500-3000 | | |

**c**

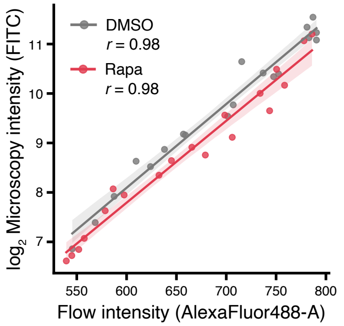


**Supplementary Figure 8 | Mycobacterial proteins are differentially degraded and sometimes re-localized with rapamycin. a** Live cell, wide-field fluorescence microscopy images of cells expressing selected target proteins with DMSO or 0.1 μg $ml^{-1}$ rapamycin. Scale bar, 5 μm. **b** Dynamic range of GFP signal for each indicated strain. **c** Correlation plot comparing fluorescent signal intensity for each tested target by microscopy and flow cytometry. Samples stratified by treatment with DMSO or 0.1 μg $ml^{-1}$ rapamycin. For **a,** data are representative images selected from among 4 fields for each and are representative of two independent experiments. In **c,** Data are bounded by the 95% confidence interval; time-lapse data represents the median fluorescent signal of all cells and flow data represents the mean fluorescent signal of two technical replicates. Related to Fig. 3.

### Supplementary Figure 9

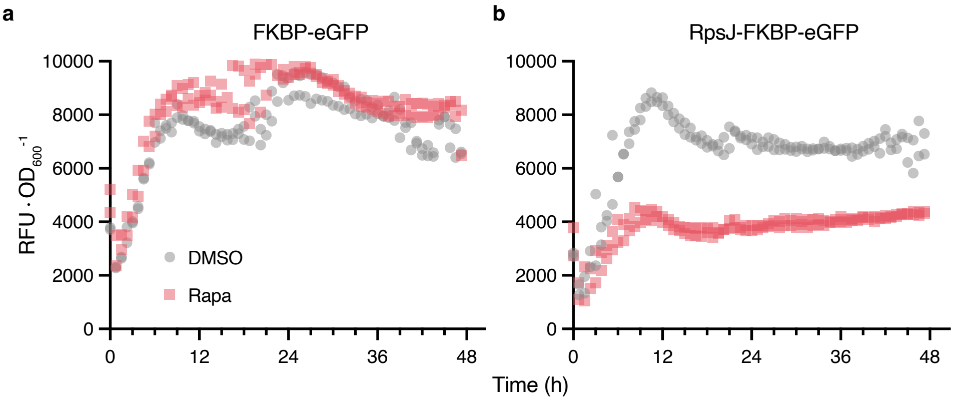


**Supplementary Figure 9 | Rapamycin stably directs degradation of RpsJ for 48 h. a-b** Cell density ($OD_{600}$)-normalized fluorescence of live cells as a proxy for protein levels of FKBP-eGFP (**a**) or RpsJ-FKBP-eGFP (**b**) in the *clpC1-frb* background. Density matched log phase cells incubated with DMSO or 1 µg $ml^{-1}$ rapamycin with shaking at 37° C. Data are individually plotted technical replicate measurements. Related to Fig. 3.

### Supplementary Figure 10

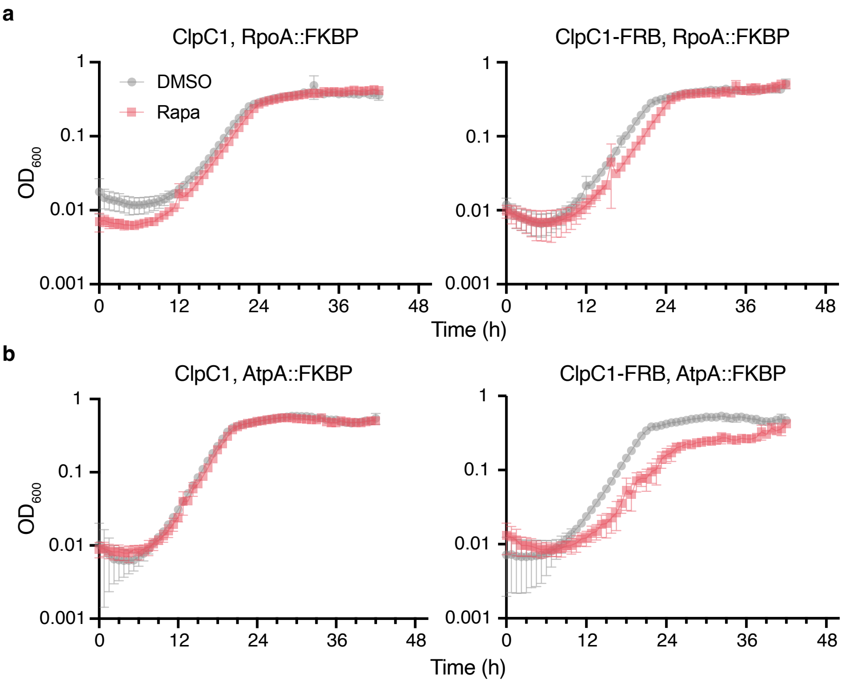


**Supplementary Figure 10 | Targeted degradation of RpoA and AtpA delays growth in liquid media. a-b** Optical density of bacterial cultures at 600nm measuring the growth kinetics of strains expressing RpoA (**a**) or AtpA (**b**) in the WT *clpC1* (left) or *clpC1-frb* (right) background when supplemented with DMSO or 10 μg ml$^{-1}$ rapamycin with shaking at 37° C. Data are mean ± s.d. of three technical replicates and are representative of two independent experiments. Related to Fig. 4.

### Supplementary Figure 11

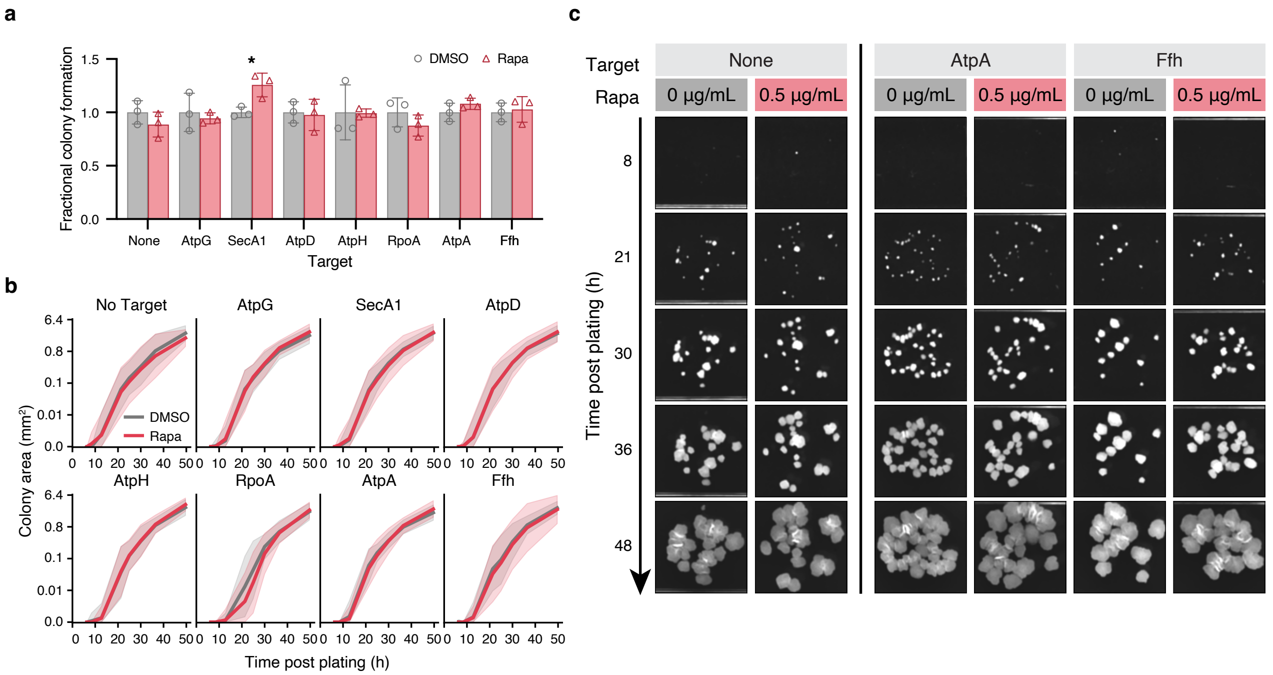


**Supplementary Figure 11 | Targeted degradation growth inhibition phenotypes are ClpC1-FRB dependent. a** Total colonies formed during outgrowth on solid media containing DMSO or 0.5 µg $ml^{-1}$ rapamycin for the indicated targets in the *clpC1* background. Density matched log phase cells serially diluted, plated on solid media containing DMSO or 0.5 µg $ml^{-1}$ rapamycin, and incubated at 37° C. *P* values were determined by unpaired two-tailed *t*-tests and compared (■) with (■). **P* < 0.05, Exact *P*-value: SecA1, ***P* = 0.0214. **b** Quantitation of colony outgrowth dynamics by colony size tracking by area ($mm^2$) of individual colonies of the indicated targets in the *clpC1* background over time. **c** Representative images illustrating colony outgrowth dynamics of the indicated targets in the *clpC1* background over time. Cells plated as in **a-b**. For **a**, data are mean ± s.d. of three technical replicates and are representative of two independent experiments; in **b**, dark lines are the mean of three technical replicates, are bounded by the 95% confidence interval, and are representative of two independent experiments. For **c**, images are representative of three technical triplicates and two independent experiments. Related to Fig. 4.

### Supplementary Figure 12

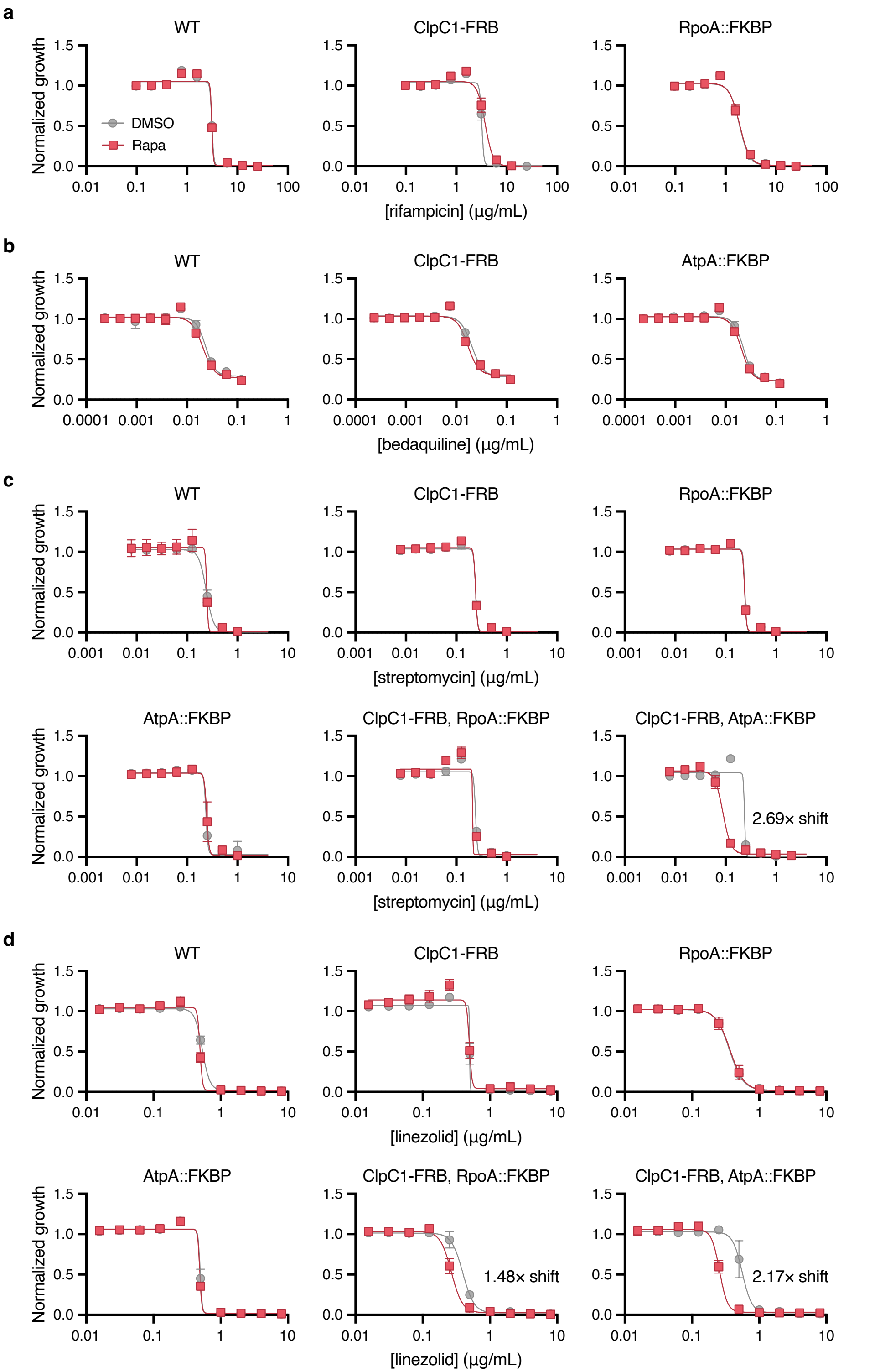


**Supplementary Figure 12 | *Msm* dose response to antibiotics with targeted protein degradation. a-d** Half-maximal minimum inhibitory concentration ($MIC_{50}$) dose response measuring the sensitivity of indicated strains to rifampicin (**a**), bedaquiline (**b**), streptomycin (**c**), or linezolid (**d**) in media supplemented with DMSO or 0.5 µg ml$^{-1}$ rapamycin. Observed fold-shifts are denoted on the corresponding plot. For **a-d**, data are mean ± s.d. of three technical replicates. Related to Fig. 5.
